# Adipocyte-Derived Amino Acid Storage Proteins are Required for Germline Stem Cell Maintenance in Adult *Drosophila* Females

**DOI:** 10.1101/2025.08.24.672014

**Authors:** Anna B. Zike, Mekenzi O. Hazen, Madison G. Abel, Eleanor B. Goldstone, Robert C. Eisman, Lesley N. Weaver

**Affiliations:** Department of Biology, Indiana University, Bloomington, IN 47405, USA

**Keywords:** Oogenesis, Fbp1, Fbp2, Lsp1, Lsp2

## Abstract

Tissue homeostasis is dependent on precise coordination between endocrine organs in response to changes in organism physiology. Secreted circulating factors from adipocytes regulate the behavior of stem cell lineages in peripheral tissues in multiple organisms. In addition to their endocrine roles, *Drosophila* adipocytes store and secrete amino acid storage proteins throughout development. During the larval feeding period, adipocytes secrete storage proteins into the hemolymph, which are reabsorbed by the adipose tissue during metamorphosis to control adult organ size and fertility. Despite the known functions for storage proteins during the larval stages, their requirement during *Drosophila* adulthood and reproduction are uncharacterized. We discover that adipocyte-specific knockdown of the storage proteins *Larval serum protein 1* (*Lsp1*) *α*/*β*/*γ* and *Larval serum protein 2* (*Lsp2*) results in a decrease in GSC maintenance. We further reveal that decreased GSC number is due to downregulation of Target of Rapamycin (TOR) signaling in GSCs, suggesting compromised amino acid sensing directly in GSCs. We also find that the proteins that mediate storage protein adipocyte reabsorption, Fat body protein 1 (Fbp1) and Fat body protein 2 (Fbp2), are expressed in ovarian follicle cells. Intriguingly, Fbp1 nor Fbp2 appear to be required in follicle cells for GSC maintenance, suggesting undiscovered requirements for amino acid storage proteins in oogenesis. Our results highlight a novel role for *Drosophila* amino acid storage proteins during adulthood and in regulating tissue stem cell lineages.

## INTRODUCTION

Multicellular organisms rely on precise coordination of circulating factors between organs to regulate developmental processes and maintain tissue homeostasis during adulthood (Castillo-Armengol et al., 2019; Droujinine and Perrimon, 2016; Katagiri, 2023), and defects in interorgan communication can result in tissue dysfunction and disease (Alvarez-Ochoa et al., 2021). For example, fasting-induced intestinal secretion of the insulin peptide (INS-7) coordinates the gut-to-brain axis to control fat mobilization in *C. elegans* (Liu et al., 2024).

Intermittent fasting in mammals activates adrenal glands to stimulate the release of free fatty acids in the hair follicle stem cell (HFSC) niche, resulting in increased levels of reactive oxygen species to induce HFSC death (Chen et al., 2025). Therefore, understanding the unique mechanisms used by different organs to communicate their physiological state can provide insight to how stem cell lineages and tissue homeostasis are maintained during adulthood.

The adult *Drosophila* ovary is an ideal model to understand interorgan control of stem cell lineages due to the conservation of signaling pathways (Hales et al., 2015; Mohr, 2018) and the unparalleled genetic toolkit that allows genetic manipulation of distinct cell types (Roote and Prokop, 2013). Each female contains two ovaries that are arranged in repeating units of 16-20 ovarioles (**Fig. 1A**) with continuously developing egg chambers. Continuous follicle production is dependent on the germline stem cell (GSC) population located in the anterior germarium (**Fig. 1B**). Each germarium houses two-to-three GSCs that divide asymmetrically to produce a self-renewing GSC and differentiating stem cell progeny (Drummond-Barbosa, 2019). Oogenesis is energetically intensive, and GSCs are sensitive to nutrient changes (Drummond-Barbosa, 2019; Drummond-Barbosa and Spradling, 2001). For example, a protein poor diet reduces GSC maintenance and results in death of differentiating progeny (Drummond-Barbosa and Spradling, 2001; Hsu and Drummond-Barbosa, 2009). Furthermore, brain-derived *Drosophila* insulin-like peptides directly regulate GSC division (LaFever and Drummond-Barbosa, 2005) and GSCs directly sense amino acids to activate Target of Rapamycin (TOR) signaling to influence GSC self-renewal and proliferation (LaFever et al., 2010). These studies suggests that complex coordination between organism physiology, organ homeostasis, and control of stem cell lineages is highly regulated in *Drosophila* females.

**Figure 1.**
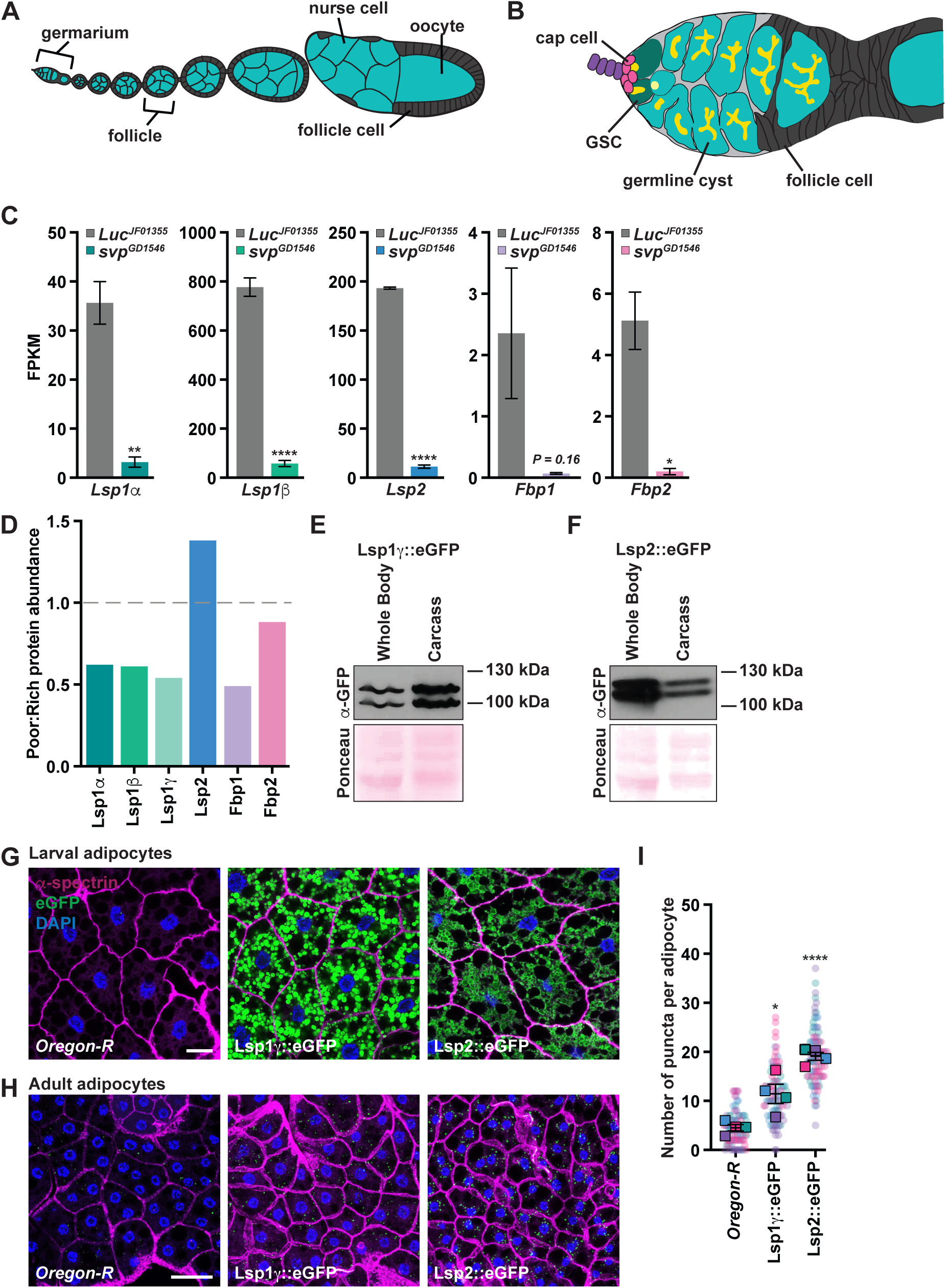
Amino acid storage proteins are present in larval and adult female fat tissue. **(A)** Cartoon schematic of the adult *Drosophila* ovariole showing the anterior germarium and progressively older egg chambers. Each egg chamber consists of germ cells (teal; 15 nurse cells and one oocyte) and somatic follicle cells (gray). **(B)** Germline stem cells (GSCs; dark teal) reside in the germarium and are maintained by a somatic niche that is composed of cap cells (pink). Germline cysts are supported by follicle cells (gray) that facilitate budding of the egg chamber. GSCs are identified by the fusome (yellow), which changes morphology as the differentiated cyst undergoes mitotic divisions with incomplete cytokinesis. **(C)** Quantification of normalized transcript expression (FPKM) for storage protein genes in control and adult fat body *svp* RNAi from (Weaver and Drummond-Barbosa, 2020). Data shown as mean ± SEM from three independent experiments. \**P*<0.05, \*\**P*<0.01, \*\*\**P*<0.001, \*\*\*\**P*<0.0001; Student’s *t*-test. **(D)** Abundance of amino acid storage proteins in adult females on a rich versus protein poor diet identified in (Matsuoka et al., 2017). **(E,F)** Western blots from whole adult females or dissected carcasses probed with anti-GFP antibodies to detect Lsp1γ::eGFP (E) or Lsp2::eGFP (F). **(G,H)** Third-instar (G) or adult (H) female adipocytes from *Oregon-R* control and amino acid storage eGFP::fusion CRISPR lines. eGFP (green), amino acid storage protein; α-spectrin (magenta), cell outline; DAPI (blue), nuclei. Scale bar = 25 µm. **(I)** SuperPlot quantifying the number of GFP puncta in *Oregon-R* control, Lsp1γ::eGFP, and Lsp2::eGFP in adult female adipocytes. Data shown as mean ± SEM from four independent samples. \**P*<0.05, \*\*\*\**P*<0.0001; Student’s *t*-test.

The adipose tissue (also known as the “fat body” in *Drosophila*) is a major site of fat storage and carbohydrate metabolism and has significant endocrine functions to mediate interorgan homeostasis through regulated secretion of adipokines (Morigny et al., 2021; Sakers et al., 2022). The fat body is composed of adipocytes and hepatocyte-like oenocytes, making it similar to the mammalian adipose and liver tissue (Arrese and Soulages, 2010; Chatterjee and Perrimon, 2021; Li et al., 2019). In *Drosophila*, adipocyte-specific metabolic activity impacts GSC maintenance. For example, loss of fatty acid beta oxidation, iron transport, and amino acid transport in adipocytes have been shown to decrease GSC number (Matsuoka et al., 2017).

Furthermore, we previously demonstrated that signaling downstream of the nuclear receptor Seven up (Svp) in adipocytes is required to maintain GSC number and prevent death of differentiating stem cell progeny (Weaver and Drummond-Barbosa, 2019). However, the precise signaling mechanisms from adipocytes to GSCs have not been fully elucidated.

In addition to lipids and fats, insect adipose tissue stores proteins in the form of amino acid storage proteins (hereafter referred to as “storage proteins”) (Haunerland, 1996; Telfer and Kunkel, 1991). Storage proteins are hexamerins composed of six identical or similar subunits that range in molecular weight from 70 to 90 kDa each (Telfer and Kunkel, 1991). Storage proteins are enriched for methionine or aromatic amino acids (phenylalanine, tryptophan, and tyrosine) and show a high degree of sequence similarity to arthropod hemanocyins (Telfer and Kunkel, 1991). In *Drosophila*, storage proteins have been extensively studied during the larval stages and metamorphosis. The *Drosophila* genome contains four storage proteins, which include the homohexamer, Larval serum protein 2 (Lsp2), and the heterohexamer composed of Larval serum protein 1α (Lsp1α), Lsp1β, and Lsp1γ. During the larval feeding period Lsp1α/β/γ and Lsp2 are synthesized by and secreted from the fat body (Burmester and Scheller, 1997; Roberts et al., 1977). The proteins encoded by *Fat body protein 1* (*Fbp1*) and *Fat body protein 2* (*Fbp2*) are the predicted receptors for LSPs (Akam et al., 1978; Wolfe et al., 1977). LSPs are reabsorbed by adipocytes during metamorphosis in a Fbp1-dependent manner, where they are utilized to control adult organ size and fitness (Valzania et al., 2024). However, Fbp1 and Fbp2 lack transmembrane domains and are found in the hemolymph (Valzania et al., 2024), suggesting an unidentified receptor may also influence storage protein reabsorption in adipocytes.

Storage proteins have also been observed in adult *Drosophila*. Larval serum protein 2 (Lsp2) was previously shown to be synthesized in the adult *Drosophila* fat body (Benes et al., 1990). While no clear role has been defined for storage proteins in adult *Drosophila*, protein levels of LSPs, Fbp1, and Fbp2 in adult female adipocytes have been shown to be sensitive to dietary changes (Matsuoka et al., 2017) and we previously reported that storage protein transcript levels are positively regulated by Svp (Weaver and Drummond-Barbosa, 2020).

Furthermore, homozygous *Drosophila Lsp1* mutants are infertile, suggesting undiscovered mechanisms for storage protein regulation of fertility in *Drosophila* (Roberts et al., 1991). These studies suggest that storage proteins may have roles in adult *Drosophila* to regulate tissue homeostasis and reproduction.

In this study we investigated the role of storage proteins in the regulation of the adult female *Drosophila* GSC lineage. Using tissue-specific manipulations, we found that adipocyte-derived storage proteins are required to regulate GSC maintenance. We found that adipocyte-derived Lsp1 and Lsp2 are required for activation of TOR signaling in GSCs for their maintenance. We also generated a suite of CRISPR knock-in fusion lines and demonstrate that storage proteins are expressed during adulthood and are also produced in the adult female ovarian follicle cells. We also present intriguing results that although Fbp1 is required for storage protein uptake in larval adipocytes, knockdown of *Fbp1* in adult female adipocytes does not influence GSC number. These results suggest a role for a yet-to-be identified intermediary processor of adipocyte-derived LSPs in maintaining GSCs. Together, our results highlight a previously unknown role for adipocyte-derived storage proteins during adulthood in the regulation of oogenesis.

## MATERIALS AND METHODS

### Drosophila strains and culture

*Drosophila* stocks were maintained on Bloomington *Drosophila* Stock Center (BDSC) Cornmeal Food that consists of 15.9 g/L inactive yeast, 9.2 g/L soy flour, 67 g/L yellow cornmeal, 5.3 g/L agar, 70.6 g/L light corn syrup, 0.059 M propionic acid at 22-25°C. Medium was supplemented with active wet yeast paste for all experiments. Flies were kept at a 12 hr/12 hr light/dark cycle at all temperatures unless otherwise noted. Previously described *Gal4* and *Gal80* transgenes were used, including *3.1Lsp2-Gal4* (Armstrong et al., 2014; Lazareva et al., 2007), *Cg-Gal4* (Asha et al., 2003), and *tubGal80^ts^*(McGuire et al., 2003). *UAS-Luc^JF01355^* (Matsuoka et al., 2017), *UAS-Fbp2^GLC01665^* (Ni et al., 2011) (Transgenic RNAi Project; <u>fgr.hms.harvard.edu/</u>), *mTOR^τιP^* (Zhang et al., 2000), *Thor^2^* (Bernal and Kimbrell, 2000), *UAS-nucGFP*, *UAS-2xEGFP*, *y^1^w^1^*, and *Oregon-R* were obtained from the Bloomington *Drosophila* Stock Center (BDSC; <u>bdsc.indiana.edu/</u>). *UAS-Fbp1^GD5143^*, *UAS-Lsp1α^KK106904^*, *Lsp1β^GD12786^*, *UAS-Lsp1γ^GD5358^*, *UAS-Lsp2^GD6088^*, and *UAS-Lsp2^KK115182^*were obtained from the Vienna *Drosophila* RNAi Stock Center (VDRC; https://shop.vbc.ac.at/vdrc_store/). Lines carrying multiple genetic elements were generated by standard crosses. Balancer chromosomes and other genetic elements are described on Flybase (www.flybase.org/).

### Generation of CRISPR Knock-In GFP-tagged transgenes

Briefly, gRNA donors were designed to knock in eGFP at the C-terminal end of each protein. Transgenesis was performed using either *yw; nosCas9 attp40* or *yw; nosCas9attp2*.

CRISPR knock-in eGFP transgenic lines were designed, microinjected, and balanced by Rainbow Transgenic Fly, Inc (Camarillo, CA). Positive eGFP transgenic lines were verified by screening for eGFP fluorescence intensity and western blot analysis.

### Western blot analysis in whole larvae and adult females

For western blot analysis, 10 mid-to-late 3^rd^ instar larvae, 10 whole females, 40 carcasses, or 40 pairs of ovaries from five-day old wet yeast fed females were collected and flash frozen in liquid nitrogen. Samples were homogenized on dry ice by hand using a tissue grinder, and 200 µl of lysis buffer [1X RIPA buffer (Fisher Scientific), 1X Halt Protease Inhibitor Cocktail (Fisher Scientific)] was added. Samples were incubated on ice for 10 minutes and centrifuged at 13,000 RPM at 4°C for 15 minutes. 150 µl of supernatant was added to 40 µl of 4X NuPAGE LDS Sample Buffer (1X, Invitrogen) plus 4% β-mercaptoethanol (Sigma Aldrich) and samples were boiled at 70°C for 10 minutes. Protein concentration was determined using the Pierce BCA Protein Assay Kit (Thermo Scientific) according to manufacturer instructions.

30 µg of total protein from each sample was loaded on a 7.5% SDS-polyacrylamide gel and transferred to nitrocellulose. Total protein was stained by incubating membranes in Ponceau (Thermo Scientific) followed by three 5-minute washes with TBST. Blots were blocked in blocking buffer [20 mM Tris pH 7.5, 150 mM NaCl (TBS), 0.1% Tween 20 (TBST), and 5% nonfat dry milk] and probed with polyclonal rabbit anti-GFP (gift from Claire Walczak, 0.25 – 1.9 µg/ml) diluted in Abdil-T (TBST, 2% BSA, and 0.1% sodium azide). Membranes were incubated overnight at 4°C, washed in TBST followed by donkey anti-rabbit IgG HRP-linked secondary antibody (1:5000 – 1:40,000) for 1 hour, washed in TBST, and detected with Clarity Western ECL Substrate Working Reagent (Bio-Rad) according to manufacturer’s instructions.

### Larval adipocyte-specific RNAi interference (RNAi)

For knockdown of eGFP-tagged storage proteins in adipocytes, 3^rd^ instar larvae carrying the *UAS-hairpin* of interest (against *Fbp1*, *Lsp1g*, or *Lsp2*) in combination with *Cg-Gal4/CyO*; *storage protein::eGFP/TM6B* (or in the case for Fbp2::eGFP, the eGFP-tagged protein was combined with the *Fbp2 RNAi* transgene and crossed with *Cg-Gal4*) were raised at 29°C to induce transgene expression during development. Amino acid storage protein knockdown larvae and their sibling controls were flash frozen and harvested for western blot analysis as described above.

### Adult tissue-specific analyses

Females containing the *UAS-transgene* of interest (*2xEGFP*, *nucGFP*, *Luc RNAi*, *Fbp1 RNAi*, *Fbp2 RNAi*, *Lsp1α RNAi*, *Lsp1β RNAi*, or *Lsp2 RNAi*) in combination with *y w; tubGal80^ts^/CyO; 3.1Lsp2-Gal4/TM3*, *y w; Thor^2^/CyO; 3.1Lsp2-Gal4 tubGal80^ts^*, *mTOR^τιP^/CyO; 3.1Lsp2-Gal4 tubGal80^ts^*, *y w; tj-Gal4 tubGal80^ts^/CyO*, or *y w; tj-Gal4 tubGal80^ts^/CyO; nSyb-Gal80/TM6B* were raised at 18°C to block Gal4 activity (and therefore, transgene expression) during development. Zero-to-2-day-old females (with *Oregon-R* males) were kept at 18°C for 2-3 days and subsequently switched to 29°C. *UAS-Luc^JF01355^*was used as an RNAi control. As additional controls, females of similar genotypes but without *Gal4/Gal80^ts^* (*UAS*-alone controls) were raised and maintained under similar conditions for 5-15 days after switching to 29°C. For all conditions, medium was supplemented with wet active yeast paste.

### Tissue immunostaining and fluorescence microscopy

Ovaries and other tissues were dissected in unsupplemented Grace’s Insect Medium (Gibco), fixed, and washed as described (Weaver and Drummond-Barbosa, 2019). Samples were blocked for at least 3 hours at room temperature in 5% normal goat serum (NGS; MP Biomedicals) plus 5% bovine serum albumin (BSA, Sigma Aldrich) in phosphate-buffered saline (PBS; 10 mM NaH_2_PO_4_/NaHPO_4_, 175 mM NaCl, pH 7.4) containing 0.1% Triton X-100 (PBST), and incubated overnight at 4°C in primary antibodies diluted in blocking solution as follows: mouse monoclonal anti-α-Spectrin (3A9) (DSHB; Developmental Studies Hybridoma Bank, 3 µg/ml); mouse monoclonal anti-Lamin C (LC28.26) (DSHB, 0.8 µg/ml); rat monoclonal anti-E-Cadherin concentrate (DCAD2-c) (DSHB, 0.37 µg/ml); rabbit monoclonal anti-p4E-BP1 (Thr37/46) (Cell Signaling Technology, 1 µg/ml), rabbit polyclonal anti-GFP (Fisher Scientific, 1 µg/ml), chicken polyclonal anti-GFP (Abcam), and rabbit polyclonal anti-Vasa (Boster Bio, 1 µg/ml). Samples were washed in PBST and incubated for 2 hours at room temperature in 1:200 Alexa Fluor 488- or 568-conjugated goat species-specific secondary antibodies (Invitrogen) in blocking solution. Stained samples were washed, mounted in Vectashield containing 1.5 µg/ml 4’,6-diamidino-2-phenylindole (DAPI) (Vector Laboratories), and imaged using a Nikon SP8 confocal microscope.

GSCs and cap cells were identified as described (Weaver and Drummond-Barbosa, 2019), and two-way ANOVA with interaction (GraphPad Prism) was used to calculate the statistical significance of any differences among genotypes in the rate of cap cell or GSC loss from at least three independent experiments, as described (Armstrong et al., 2014).

For Dad::nlsGFP quantification, the densitometric mean of individual GSC nuclei was measured from optical sections containing the largest nuclear diameter (visualized by DAPI) using FIJI (https://fiji.sc/) (Armstrong et al., 2014). For E-Cadherin quantification, the total densitometric value from maximum intensity projections around the cap cells (identified using LamC staining) was measured with FIJI (Weaver and Drummond-Barbosa, 2018).

Quantification of E-Cadherin levels by immunofluorescence is a well-established method to determine whether adherence of GSCs to the niche is compromised (Hsu and Drummond-Barbosa, 2011). p4E-BP1 levels were determined by taking Z-stacks through the whole GSC and total stack intensity was projected. Densitometric data for Dad::nlsGFP, E-Cadherin, and p4E-BP1 levels were subjected to the Mann-Whitney *U*-test.

For lipid droplet visualization, fixed and washed fat bodies attached to abdominal carcasses were incubated with 2.5X Alexa Fluor 488-conjugated phalloidin (Molecular Probes) in PBST for 20 min, rinsed, and washed three times for 15 min each in PBST. Abdominal carcasses were incubated in 25 ng/ml Nile Red dye for 10 min. Samples were stored in Vectashield plus DAPI (Vector Laboratories) and fat bodies were scraped off the abdominal carcass and imaged using a Nikon SP8 confocal. The largest cell area of each adipocyte (based on phalloidin staining) used to measure adipocyte area; whereas the longest width of each nucleus per adipocyte (based on DAPI) determined nuclear diameter. Each measurement was performed using FIJI. Three independent experiments were performed, and statistical analysis was performed using a Student’s *t* test.

For analysis of eGFP-tagged amino acid storage protein localization in female adipocytes, larval fat bodies were fixed in 5.3% formaldehyde in Grace’s media for 13 min (20 min for adult female fat) at room temperature. Samples were subsequently rinsed and washed three times for 15 min each in 1X PBS containing 0.5% Tween. Samples were blocked for at least 3 hours in block solution containing 1X PBS, 0.5% Tween, 5% NGS, and 5% BSA. Samples were incubated in polyclonal rabbit anti-GFP (Fisher Scientific, 1 µg/ml) at 4 degrees overnight. Fat bodies were washed in PBS-0.5% Tween and incubated for 2 hours at room temperature in 1:200 Alexa Fluor 488-conjugated goat anti-rabbit secondary antibody (Invitrogen) in blocking solution. Samples were subsequently incubated in 2.5X Alexa-594 phalloidin (Molecular Probes) in PBS-0.5% Tween for 20 min, rinsed, and washed three times for 15 min each in PBS-0.5% Tween. Stained samples were washed, mounted in Vectashield containing 1.5 µg/ml 4’,6-diamidino-2-phenylindole (DAPI) (Vector Laboratories). Images were collected on a Nikon SP8 confocal microscope at 0.5 µm intervals, and 3-5 z-planes were combined for analysis. For adult fat eGFP analysis, puncta from at least 25 adipocytes were measured from at least four independent experiments, and statistical analysis was performed using a paired Student’s *t* test.

### ApopTag Assays

To detect dying germline cysts, the ApopTag Indirect *In Situ* Apoptosis Detection Kit (Millipore Sigma) was used according to the manufacturer’s instructions as previously described (Weaver and Drummond-Barbosa, 2019). Briefly, fixed and teased ovaries were rinsed in equilibration buffer twice for 5 minutes each at room temperature. Samples were incubated in 100 µl TdT solution at 37°C for 1 hour with mixing at 15-minute intervals. Ovaries were washed three times in 1X PBS followed by incubation in anti-digoxigenin conjugate for 30 min at room temperature protected from light. Samples were washed four times in 1X PBS and processed for immunofluorescence as described above.

### RNA isolation

Abdominal carcasses from 100 females of each genotype were dissected in Grace’s medium supplemented with 10% fetal bovine serum (FBS; Sigma) as previously described (Weaver and Drummond-Barbosa, 2020). Fat body cells were dissociated from abdominal carcasses with 500 µl dissociation buffer [0.2% Trypsin (Sigma) plus 2 mg/ml collagenase (Sigma) in 1x PBS] per 50 carcasses for 30 minutes at room temperature. Samples were gently agitated every 5 minutes to facilitate separation of cells from the cuticle. 500 µl of Grace’s media plus 10% FBS was added to stop the enzymatic reaction, and supernatants were placed in new tubes. Carcasses were rinsed with Grace’s medium plus 10% FBS and the two supernatants per genotype were combined. Dissociated cells were centrifuged at 3.3 rpm for 5 min at room temperature. Supernatants were removed and cells were flash frozen in liquid nitrogen. Cells were lysed in 250 µl lysis buffer from the RNAqueous-4PCR DNA-free RNA isolation for RT-PCR kit (Ambion). RNA was extracted from all samples following the manufacturer’s instructions. Three independent experiments were performed for quantitative reverse-transcriptase polymerase chain reaction (qRT-PCR) analysis.

### cDNA synthesis and quantitative reverse-transcriptase polymerase chain reaction (qRT-PCR)

cDNA was synthesized from 500 ng of total RNA described above for each sample using Superscript II Reverse Transcriptase (Thermo Fisher Scientific) according to the manufacturer’s instructions. PowerUp SYBR Green Master Mix (Thermo Fisher Scientific) was used for RT-qPCR. The reactions for three independent biological replicates were performed in triplicate using LightCycler 96 (Roche). Amplification fluorescence threshold was determined by LightCycler 96 software, and ddCT were calculated using Microsoft Excel. Fold change of transcript levels was calculated in Excel as described (Taylor et al., 2019). The primers used for all PCR reactions are listed in Table S1. *Rp49* and *Act5C* transcript levels were used as references.

### Survival assays

For survival analysis, four-to-six vials containing 25 two-to-four-day old, mated females were incubated at 29°C. Vials were flipped every 1–3 days until no living flies remained in the vials. Death was recorded when the flies were flipped to fresh food. Data was plotted in GraphPad Prism (GraphPad) and survival curve statistical analysis was performed using a log-rank test.

### Egg count assays

Egg count analysis was performed as previously described (Weaver and Drummond-Barbosa, 2019). Briefly, five females of the appropriate genotype and five *y^1^ w^1^* males were maintained at 29°C in perforated plastic bottles capped with molasses/agar plates supplemented with active yeast paste. Plates were changed twice daily, and the number of eggs per day was counted in five replicates per genotype. Data was subjected to a paired Student’s *t*-test.

### Fly Liquid-Food Interaction Counter (FLIC) assay

The Fly Liquid-Food Interaction Counter (FLIC) system was used to determine differences in feeding behavior as previously described (Fleck et al., 2024; Ro et al., 2014). FLIC *Drosophila* Feeding Monitors (DFMs, Sable Systems International, models DFMV2 and DFMV3) were used in the single choice configuration and each chamber was loaded with liquid food solution [4% sucrose (m/v), 1.5% yeast extract (m/v)]. Females incubated at 29°C for 10 days were switched from solid food to liquid food at 29°C overnight (∼8-12 hours) prior to running each experiment. Flies were aspirated into the DFM chambers and feeding behavior was measured for 24 hours. Each FLIC experiment contains pooled data from at least 30 flies for each genotype. FLIC data were analyzed using previously described custom R code (Ro et al., 2014), which is available at https://github.com/PletcherLab/FLIC_R_Code. Default thresholds were used for analysis except for the following: minimum feeding threshold = 10, tasting threshold = (0,10). Animals that did not participate (i.e., returned zero values), whose DFM returned an unstable baseline signal, or who produced extreme outliers (i.e., exceeding twice the mean of the population) were excluded from analysis. Data were subjected to a Mann-Whitney *U*-test.

## RESULTS

### Adipocyte-derived amino acid storage proteins are required for GSC maintenance

Storage proteins are required for fecundity in many adult insects; however, the requirement for storage proteins during oogenesis in *Drosophila* has been understudied. We previously performed transcriptomics on the adult female fat body to identify signaling components downstream of Svp. Analysis of this data revealed that *Lsp1α*/*β*, *Lsp2*, and *Fbp2* transcripts were significantly downregulated in females where *svp* was depleted from the fat body (**Fig. 1C**; (Weaver and Drummond-Barbosa, 2020); however, it is unknown whether LSPs or Fbp2 are direct targets of Svp from this dataset. Furthermore, adipocyte-specific affinity labeling of secreted peptides identified storage proteins in adult hemolymph (Droujinine et al., 2021), and a previous proteomics analysis determined that storage proteins are sensitive to changes in diet in adult female adipocytes [**Fig. 1D**; (Matsuoka et al., 2017)]. We therefore decided to test the involvement of adipocyte-derived storage proteins in regulating the GSC lineage.

To assess whether storage proteins are expressed in adult females, we utilized a CRISPR knock-in strategy to express enhanced GFP (eGFP) at the C-terminal end of the endogenous protein (Lsp1γ and Lsp2) and analyzed protein expression in female larval and adult adipocytes by western blot analysis (**Fig. 1E and F; Fig. S1A and B**). To test the specificity of our eGFP knock-in lines, we knocked down *Lsp1*γ and *Lsp2* in larval adipocytes during development using the *Cg-Gal4* driver (Asha et al., 2003) and observed a significant decrease in eGFP-tagged protein levels relative to controls (**Fig. S1A and B**). We next analyzed the expression pattern in adipocytes by immunofluorescence using *Oregon-R* as a negative control (**Fig. 1G and H**). In larval female adipocytes, Lsp1γ::eGFP (representing the heterohexameric complex of Lsp1α/β/γ) and Lsp2::eGFP are abundantly expressed (**Fig. 1G**) and form granule-like structures in the cytosol. In contrast, storage protein expression in adult female adipocytes is significantly decreased compared to larval female adipocytes (**Fig. 1H**), showing small Lsp1γ::eGFP and Lsp2::eGFP puncta (**Fig. 1I)**. Collectively, these results suggest that although storage proteins are not as highly expressed in the adipocytes of adult females compared to larvae, they are detected and may have functional roles during adulthood.

### Adipocyte-derived amino acid storage proteins are required for GSC maintenance

We next asked whether adipocyte-secreted Lsp1α/β/γ and Lsp2 are required for early processes of oogenesis (GSC maintenance or survival of early germline cysts). Individual knockdown of *Lsp1α* or *Lsp1β* is sufficient to decrease the transcript levels of all components of the Lsp1 complex (**Fig. S2A, B, and D**; (Valzania et al., 2024). Therefore, we knocked down *Lsp1α*, *Lsp1β*, or *Lsp2* (**Fig. S2C and E**) specifically in adult adipocytes using the previously described *tubGal80^ts^; 3.1Lsp2-Gal4* (*3.1Lsp2^ts^*) driver (Armstrong et al., 2014) and assayed GSC number over 15 days by staining ovaries with α-Spectrin to label GSCs and Laminin C to label cap cell nuclear lamina (**Fig. 2A and B**). Knockdown of *Lsp1α/β* and *Lsp2* in adult female adipocytes significantly increased the rate of GSC loss compared to *Luc* control (**Fig. 2C and D; Fig. S2F and G**). The loss of GSCs was not due to leaky expression of the *UAS* transgenes, as there was no significant decrease in GSC number of *UAS-alone* controls over time compared to the *UAS-Luc* RNAi hairpin (**Fig. S2H and I**). In addition, the decrease in GSC maintenance was not dependent on the size of the niche (Hsu and Drummond-Barbosa, 2009), as cap cell numbers were unaffected (**Fig. S2J and K**). Furthermore, GSC loss was not due to gross disruption of adipose tissue as measured by adipocyte nuclear or cell size (**Fig. S3**).

**Figure 2.**
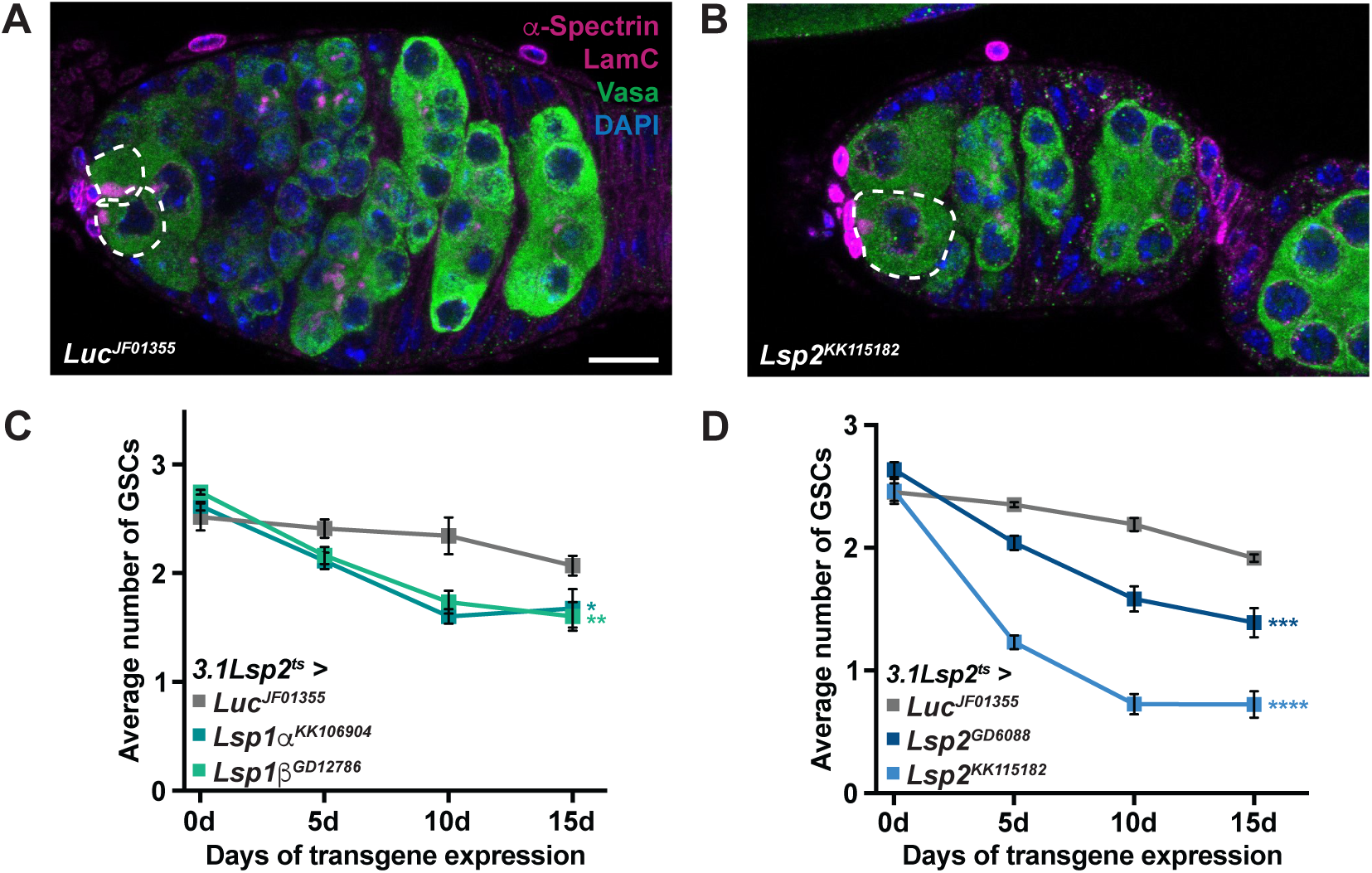
Adult adipocyte-specific knockdown of larval serum proteins decreases GSC maintenance. (A,B) Germaria from *Luc* control (A) and adipocyte-specific *Lsp1α* knockdown (B) after 15 days of transgene expression. Vasa (green), germ cells; α-spectrin (magenta), fusome; LamC (magenta), cap cell nuclear lamina; DAPI (blue), nuclei. GSCs are outlined. Scale bar, 10 µm. **(C,D)** Average number of GSCs over time in *Luc* control females and from adipocyte-specific *Lsp1α/β* (C) or *Lsp2* (D) knockdown. Data shown as mean ± standard error of the mean (SEM) from three independent experiments. \**P*<0.05, \*\**P*<0.01, \*\*\**P*<0.001, \*\*\*\**P*<0.0001; two-way ANOVA with interaction.

It was recently shown that dietary restriction due to reduction of *Lsp2* in larvae increases lifespan during adulthood (Kosakamoto et al., 2025). Consistent this, we also observed a slight, but significant lifespan enhancement when *Lsp1a* or *Lsp2* were knocked down specifically in adult adipocytes (**Fig. S4A and B**), suggesting that dietary restriction (via lower storage protein levels) may enhance lifespan at the expense of reproductive fitness. We also explored whether loss of LSPs in adipocytes altered feeding behavior using the Fly Liquid food Interaction Counter (FLIC) and did not observe any change in the number of feeding events or duration of feeding compared to *Luc* RNAi control (**Fig. S4C-F**).

Notably, storage protein knockdown did not influence the number of eggs laid by females compared to *Luc* control over a two-week period (**Fig. S4G**). It should be noted, however, that egg count analyses do not always reveal subtle changes in oogenesis (Ma et al., 2020; Weaver and Drummond-Barbosa, 2019), especially defects occurring at early stages in the germline. Production of a mature oocyte from a GSC takes approximately seven days (He et al., 2011) and changes in GSC maintenance due to adipocyte-specific storage protein knockdown are not observed until 10 days after transgene expression; therefore, it is unlikely that changes in egg production would be observed by 15 days. Furthermore, egg production is significantly inhibited at higher temperatures in wild-type strains (Gandara and Drummond-Barbosa, 2022), preventing long-term observation of consequences from GSC maintenance defects. Our data suggest that although loss of *Lsp1a* and *Lsp2* enhances adult female lifespan, additional physiological effects that could influence GSC maintenance were unaffected.

To assess whether knockdown of *Lsp1α*/*β*/*γ* or *Lsp2* influences survival of early germline cysts, we labeled germaria with ApopTag, which is a TUNEL-based assay to detect significant DNA fragmentation that is indicative of apoptosis (Drummond-Barbosa and Spradling, 2001).

Compared to *Luc* control, knockdown of *Lsp1α/β/γ* or *Lsp2* in adipocytes did not significantly increase the percentage of ApopTag positive germline cysts 10 days after transgene expression (**Fig. S5**). Collectively, these results suggest that Lsp1α/β/γ and Lsp2 are required in adult female adipocytes to control GSC maintenance.

### Storage protein reabsorption by Fbp1 or Fbp2 does not regulate GSC maintenance

Reabsorption of storage proteins in adipocytes during pupariation is dependent on Fbp1 (Valzania et al., 2024). Considering Fbp1 and Fbp2 proteins are present in adult female fat and are also regulated by a protein-rich diet, we hypothesized that Fbp1-mediated retrieval of storage proteins may promote GSC self-renewal. We assessed Fbp1 and Fbp2 expression in adult female adipocytes using our eGFP CRISPR knock-in constructs, which were confirmed to be specific based on western blot analysis of knockdown of *Fbp1* or *Fbp2* in the larval fat body compared to control (**Fig. S1C and D**).

Immunofluorescence analysis showed that Fbp1::eGFP is expressed in granules in female larvae adipocytes, however Fbp2::eGFP has a more diffuse expression pattern (**Fig. 3A**). In contrast, Fbp1 and Fbp2 expression in adult female adipocytes is decreased and shows variable puncta levels compared to the *Oregon-R* control (**Fig. 3B and C**). Given the lower amounts of Fbp1 and Fbp2 in adult adipocytes compared to larvae, we confirmed Fbp1 and Fbp2 expression in adult female fat bodies by western blot (**Fig. 3D and E**). Fbp1 is an ecdysone-inducible gene and is cleaved to generate a 64 kDa fragment (Burmester et al., 1999). Fbp1 is further cleaved when reabsorbed by adipocytes to generate a 47 kDa fragment (Burmester et al., 1999). Taking into consideration the eGFP tag (∼27 kDa), the multiple bands present in the Fbp1::eGFP western blots are consistent with ecdysone-induced cleavage of Fbp1.

**Figure 3.**
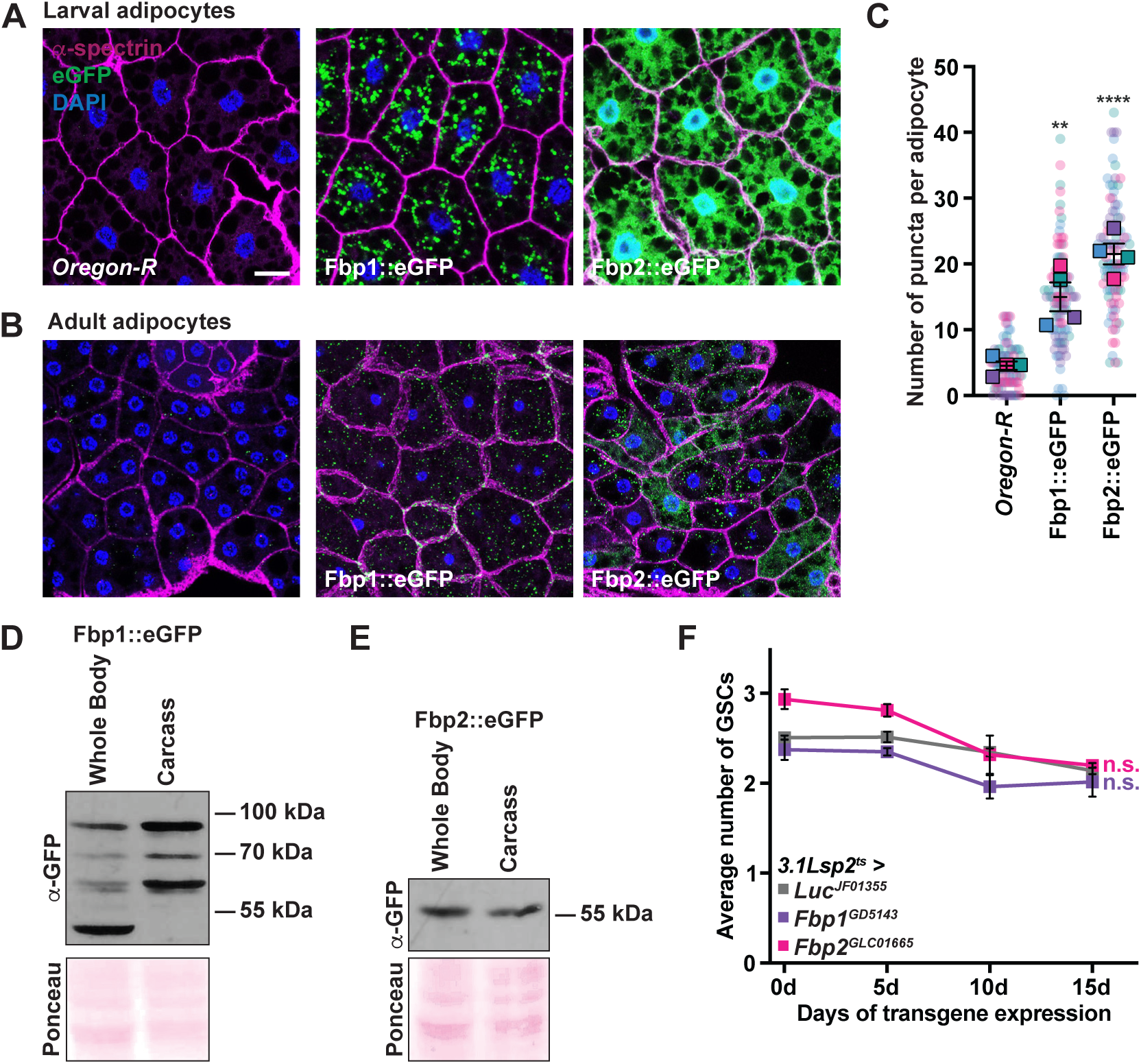
Knockdown *Fbp1* or *Fbp2* in adult female adipocytes does not influence GSC number. (A,B) Third-instar (A) or adult (B) female adipocytes from *Oregon-R* control and Fat body protein eGFP::fusion CRISPR lines. eGFP (green), amino acid storage p**rotein; α-spectrin (magenta), cell outline; DAPI (blue), nuclei. Scale bar = 25 µm. Note that the *Oregon-R* control is identical to Figure 1G and H** as all of the fat samples were collected and analyzed at the same time and under the same conditions. **(C)** SuperPlot showing quantification of the number of GFP puncta in *Oregon-R* control, Fbp1::eGFP, and Fbp2::eGFP in adult female adipocytes. Data shown as mean ± SEM from four independent samples. \*\**P*<0.01, \*\*\*\**P*<0.0001; Student’s *t*-test. **(D,E)** Western blots from whole adult females or dissected carcasses probed with anti-GFP antibodies to detect Fbp1::eGFP (D) or Fbp2::eGFP (E). **(F)** Average number of GSCs over time for adipocyte-specific *Fbp1* or *Fbp2* knockdown compared to *Luc* control. Data shown as mean ± SEM. n.s., no significant differences; two-way ANOVA with interaction.

To determine whether receptor-mediated adipocyte reabsorption of storage proteins is required for regulation of GSC number, we knocked down *Fbp1* (as well as the other predicted storage protein receptor, *Fbp2*) in adult female adipocytes (**Fig. S6A-C**) and analyzed GSC maintenance. Relative to *Luc* control, knockdown of *Fbp1* or *Fbp2* did not significantly decrease GSC maintenance over time (**Fig. 3F; Fig. S6D**). These results suggest that Fbp1 nor Fbp2 are required in adult female adipocytes for maintaining GSC number.

### Fbp1 and Fbp2 are expressed in ovarian follicle cells but are not required for GSC maintenance

Because Fbp1 and Fbp2 are not required in adult female adipocytes for regulation of GSC number, we hypothesized that Fbp1 or Fbp2 may be required in the ovarian germline or somatic cell populations for storage protein absorption. We therefore assayed ovarian expression of our CRISPR eGFP knock-in lines for expression in the ovary. Lsp1γ::eGFP was not detected in either the germline or the soma of the adult ovary by immunofluorescence or western blot (**Fig. S7A and B**). Interestingly, we did detect Lsp2::eGFP in adult ovaries (**Fig. S7C**), and immunofluorescence showed Lsp2::eGFP in the oocyte of egg chambers as early as stage 9 (**Fig. S7D**). This suggests Lsp2 may be synthesized in oocytes or possibly transported to the oocyte from another tissue (e.g., adipocytes) since *Lsp2* transcripts are not detected by single cell sequencing of the ovary (Li et al., 2022; Rust et al., 2020). Notably, both Fbp1::eGFP (**Fig. 4A**) and Fbp2::eGFP (**Fig. 4B**) are detected in the adult ovary by western blot.

**Figure 4.**
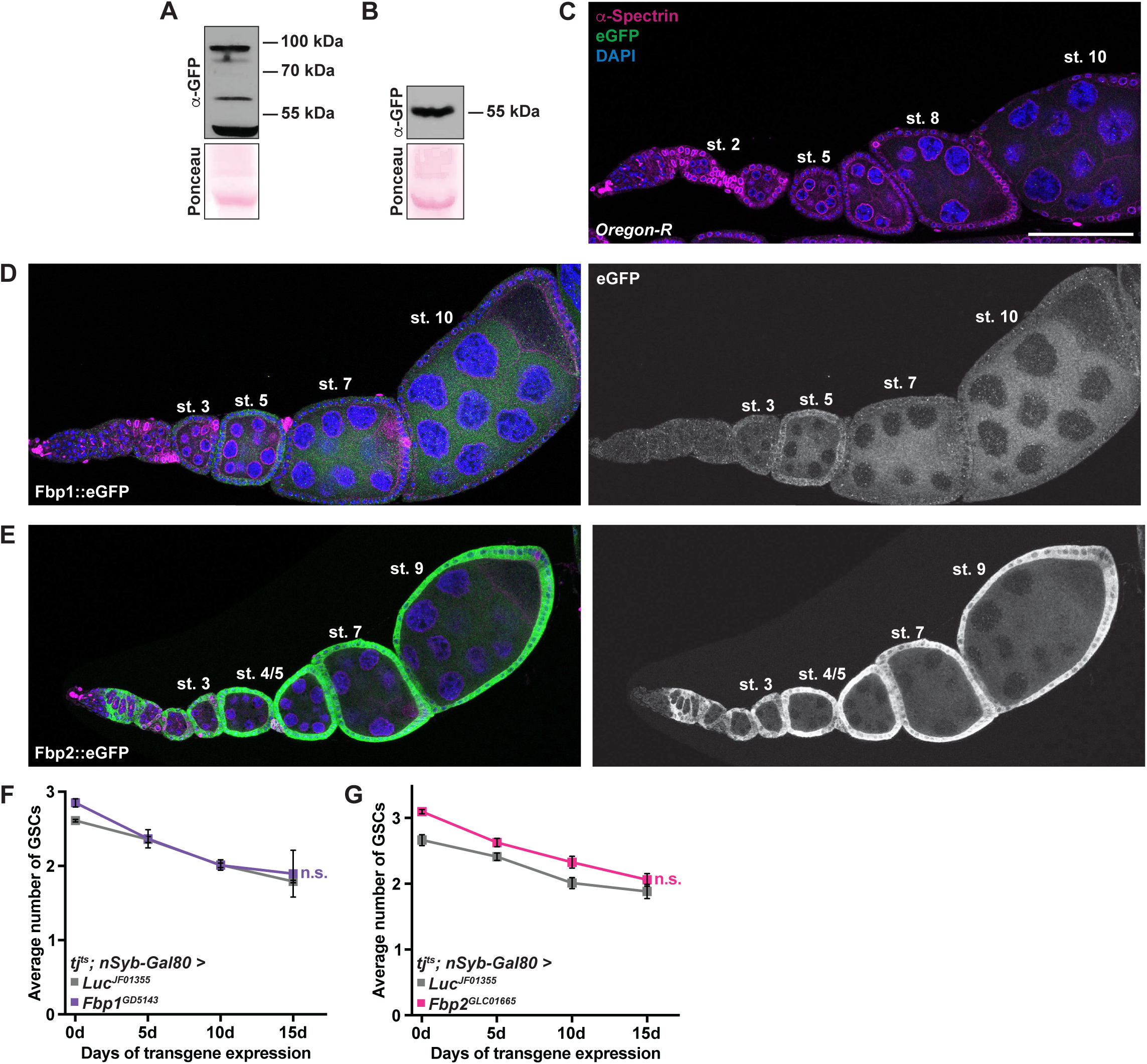
Fbp1 and Fbp2 are expressed in adult ovarian follicle cells, but are not required for GSC maintenance. (A,B) Western blot analysis from whole adult females and carcasses probed with anti-GFP antibody to detect Fbp1::eGFP (A) or Fbp2::eGFP (B). **(C-E)** Ovarioles from *Oregon-R* (C), *Fbp1::eGFP* (D), and *Fbp2::eGFP* (E) adult females. eGFP (green), amino acid storage protein; α-spectrin (magenta), fusome; LamC (magenta), cap cell nuclear lamina; DAPI (blue), nuclei. Scale bar = 100 µm. **(F,G)** Average number of GSCs over 15 days from follicle cell knockdown of *Fbp1* (F) and *Fbp2* (G). Data shown as mean ± SEM from three independent experiments. n.s., no significant differences; two-way ANOVA with interaction.

Immunofluorescence analysis showed that relative to *Oregon-R* control (**Fig. 4C**), Fbp1::eGFP localized to ovarian follicle cells starting in stage 3 egg chambers and decreased expression after stage 5 (**Fig. 4D**). Fbp1::eGFP was also detected in the cytoplasm of nurse cells beginning in stage 7 egg chambers. In contrast, Fbp2::eGFP exclusively localized to follicle cells (**Fig. 4E**), beginning in the germarium and continuing through the remainder of oogenesis. To our knowledge, this is the first reported localization of Lsp2, Fbp1, and Fbp2 in the adult *Drosophila* ovary, suggesting functional roles in adult females.

To determine whether Fbp1 or Fbp2 are required in ovarian follicle cells for GSC number, we designed a modified transgenic line of the commonly used follicle cell driver *tj-Gal4* (Sahai-Hernandez and Nystul, 2013). In addition to follicle cell expression, *tj-Gal4* has ectopic expression in the central nervous system, Malpighian tubules, and adipocytes (**Fig. S7E**; (Weaver et al., 2020). To alleviate expression in the brain, we combined the temperature inducible *tj-Gal4 tubGal80^ts^*driver with *nSyb-Gal80* (to prevent neuronal Gal4 expression independent of temperature), which significantly decreased expression in the brain when females were incubated at 29°C (**Fig. S7F**). Note there is still significant expression in adipocytes; however, because Fbp1 and Fbp2 are not required in adipocytes to regulate GSC number, we reasoned any differences in GSC maintenance would be due primarily to their requirement in follicle cells. However, knockdown of *Fbp1* or *Fbp2* using *tj-Gal4 tubGal80^ts^; nSyb-Gal80* did not significantly decrease the number of GSCs over 15 days relative to *Luc* control (**Fig. 4F and G; Fig. S7G and H**). These results suggest that although Fbp1 and Fbp2 are expressed in the follicle cells of the ovary, they are not required for GSC maintenance.

### Adipocyte-derived amino acid storage proteins do not influence BMP signaling and E-Cadherin levels in GSCs

GSCs are maintained in the niche primarily through Bone Morphogenic Protein (BMP) signaling from cap cells (Song and Xie, 2002) and by E-Cadherin adhesion to cap cells (Song et al., 2004) to prevent premature differentiation. We asked whether loss of GSCs due to knockdown of *Lsp1*α/β/γ and *Lsp2* in adult adipocytes was mediated by decreased BMP signaling or a reduction in E-Cadherin levels at the niche. To test whether BMP signaling was affected, we measured the nuclear intensity of the Dad::nlsGFP BMP signaling reporter (Ayyaz et al., 2015) when *Lsp1*α/β or *Lsp2* were knocked down specifically in adult female adipocytes after 10 days. Storage protein loss specifically in adult female adipocytes did not have a significant decrease on Dad::nlsGFP levels relative to *Luc* control (**Fig. S8A-C**), suggesting that adipocyte-derived amino acid storage proteins do not regulate BMP signaling. Furthermore, adipocyte-specific loss of storage proteins did not affect E-Cadherin levels in the GSC-niche interface, suggesting a different mechanism is required to maintain GSCs (**Fig. S8D-F**).

### Adipocyte-derived storage proteins regulate GSC Target of Rapamycin signaling

mTOR is a conserved kinase that regulates cell growth and survival downstream of amino acids (Gonzalez and Hall, 2017) and is required in adult female GSCs for stem cell maintenance (LaFever et al., 2010). We hypothesized that adipocyte-derived storage proteins may be transported to the ovary (or some other peripheral tissue) and are catabolized into amino acids to promote TOR signaling. To test this hypothesis, we knocked down *Lsp1α/β* or *Lsp2* specifically in adult female adipocytes using *3.1Lsp2-Gal4^ts^* and assayed the levels of phosphorylated 4E-BP1 (p4E-BP1), the eukaryotic translation initiation factor 4E binding protein that is phosphorylated by mTOR, which serves as a direct read-out of TOR activity (**Fig. 5A**; (LaFever et al., 2010; Wei et al., 2019). In female GSCs, p4E-BP1 is present during M phase but not G phase (LaFever et al., 2010), which limited the number of GSCs that could be analyzed since mitotic pHH3+ GSCs are only observed in about 5% of the population (Hinnant et al., 2017). Regardless, adipocyte-specific knockdown of amino acid storage proteins significantly decreased p4E-BP1 levels (**Fig. 5B**). Notably, there was no difference in p4E-BP1 levels at different mitotic phases (e.g., prometaphase versus metaphase) within an RNAi condition (**Fig. S9A and B**), suggesting the phosphorylation status of 4E-BP1 does not change during early phases of mitosis.

**Figure 5.**
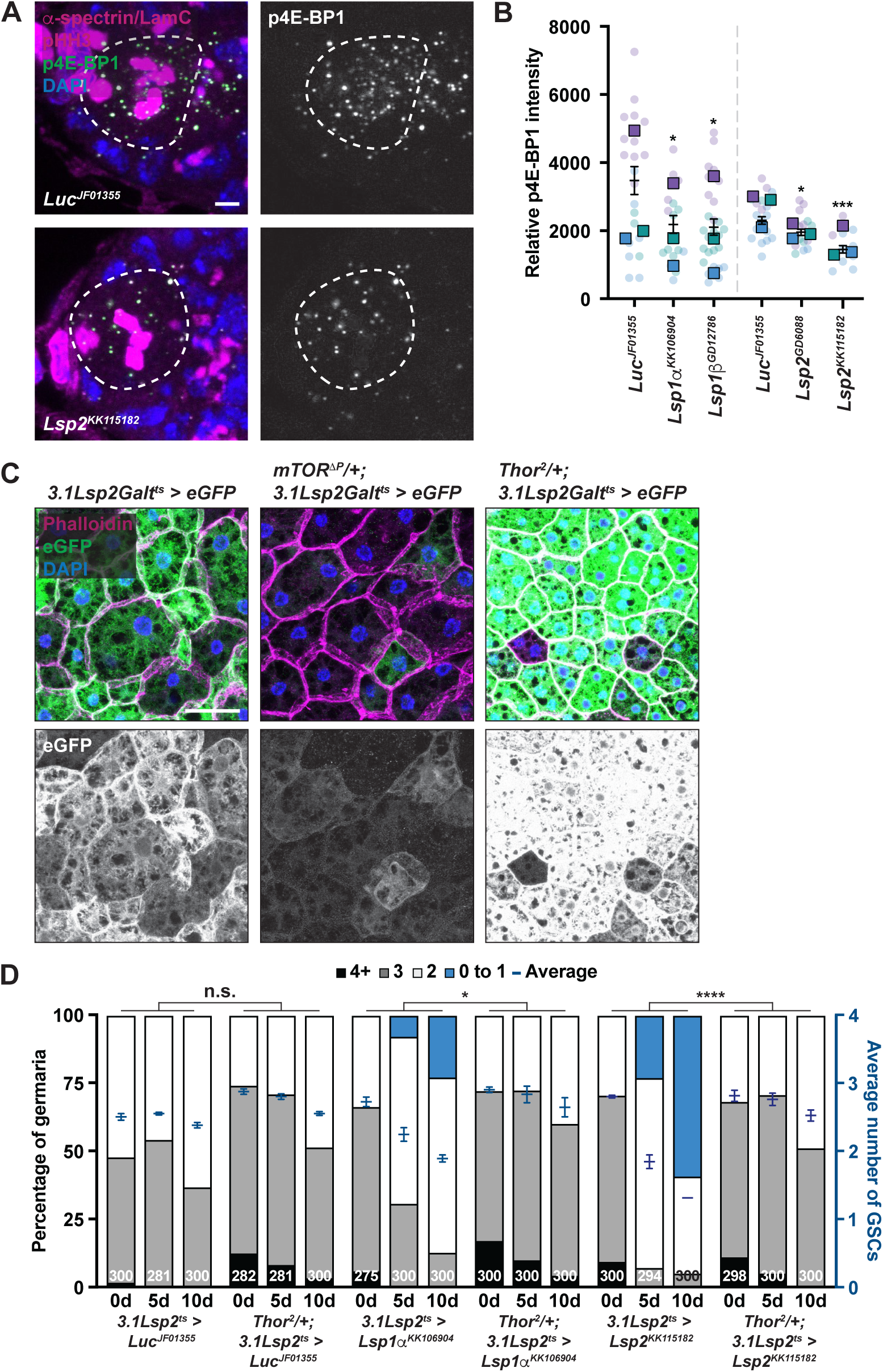
Adipocyte-specific knockdown of storage proteins decreases TOR signaling in GSCs. **(A)** Anterior portion of germaria from adipocyte-specific *Luc* control and *Lsp2* knockdown 10 days after transgene induction. Phosphorylated 4E-BP1 (p4E-BP1, green), a direct readout of mTOR signaling (LaFever et al., 2010; Wei et al., 2019); α-spectrin (magenta), fusome; LamC (magenta), cap cell nuclear lamina; DAPI (blue), nuclei. Scale bar = 2.5 µm. The dotted lines outline GSCs. **(B)** SuperPlot of p4E-BP1 intensity per GSC for each genotype. At least 20 GSCs per genotype were analyzed from three independent experiments. Data shown as mean ± 95% confidence interval for each genotype. \**P* < 0.05, \*\*\*\**P* < 0.0001; Mann-Whitney *U*-test. **(C)** Representative images of adult female adipocytes from *3.1Lsp2-Gal4^ts^*, *mTOR^DP^/+; Lsp2-Gal4^ts^*, or *Thor^2^/+; Lsp2-Gal4^ts^* expressing *UAS-2xEGFP*. EGFP (green), Phalloidin (cell membrane; magenta), DAPI (blue), nuclei. Scale bar = 25 µm. **(D)** Frequencies of germaria containing zero-or-one, two, three, or four-or-more GSCs at multiple times points of adipocyte-specific RNAi against *Luc* (control), *Lsp1a*, or *Lsp2* in females carrying one copy of *Thor^2^* compared to their wild-type controls. The average number of GSCs per germarium is plotted in dark blue. The number of germaria analyzed is shown inside the bars. Data shown as mean ± SEM. \**P* < 0.05, \*\*\*\**P* < 0.0001; two-way ANOVA with interaction.

To further test whether adipocyte-derived storage proteins regulate mTOR signaling in GSCs, we examined genetic interactions between adipocyte-derived LSPs, *mTOR*, and *Thor* (4E-BP1 in *Drosophila*, a negative regulator of TOR signaling). First, we reasoned that if adipocyte LSPs regulate GSC maintenance though TOR signaling, decreasing the levels of *mTOR* would enhance the GSC loss phenotype, whereas decreasing *Thor* levels would rescue GSC loss. We first combined the *mTOR^τιP^* mutant allele [a P-element insertion that that removes the translation start site and first 902 codons (Zhang et al., 2000)] with *3.1Lsp2^ts^*. RT-qPCR analysis confirmed a significant reduction in *mTOR* transcripts (**Fig. S9C**). However, the *3.1Lsp2-Gal4* driver in a heterozygous *mTOR^τιP^* background displayed less robust expression compared to control (**Fig. 5C**), preventing further analysis.

We next generated a line combining *3.1Lsp2-Gal4^ts^* with the *Thor^2^* mutant [an amorphic allele generated from an imprecise P-element excision that removes the entire coding sequence (Bernal and Kimbrell, 2000)]. In this line, *Thor* mRNA was significantly reduced compared to control (**Fig. S9C**) and driver expression was increased (**Fig. 5C**). GSC maintenance was unaffected by one copy of *Thor^2^* in *Luc* control conditions (**Fig. 5D**, compare control and experimental slopes). By contrast, GSC maintenance due to adipocyte loss of LSPs was significantly rescued by one copy of *Thor^2^*. Collectively, our results suggest that adipocyte-derived storage proteins act upstream of TOR in GSCs to control GSC maintenance.

## DISCUSSION

Organism physiology is dependent on modulation of interorgan communication in response to changes in macromolecule levels. Diet-derived amino acids and amino acid sensing are required in multiple cell types to promote stem cell proliferation and self-renewal to maintain tissue homeostasis. In insects, amino acid storage proteins are essential for metamorphosis and organ growth, as well as reproduction. Our results herein show that adipocyte-derived storage proteins are required in adult *Drosophila* females to regulate GSC maintenance to facilitate egg production. Furthermore, our results highlight an uncharacterized requirement for storage proteins in adulthood for female reproduction in *Drosophila*.

### Storage proteins are expressed in adult insects

Although the requirement of storage proteins has been primarily studied for their roles during larval feeding and metamorphosis, storage protein production has also been observed in adult insects (Benes et al., 1990; Martins et al., 2008; Wheeler and Buck, 1995, 1996; Yan et al., 2022). For example, transcripts of the honeybee storage protein *hex 70a* are more abundant in adult female workers compared to males, and there is a significant increase in *hex 70a* fat body transcripts in workers with active ovaries compared to those with inactive ovaries (Martins et al., 2008). Furthermore, *Lsp2* transcripts are detected in the adult *Drosophila* fat body, and the protein accumulates in the hemolymph, albeit at levels significantly decreased compared to larvae (Benes et al., 1990). These results are consistent with our detection of storage proteins (Lsp2::GFP, Fbp1::GFP, and Fbp2::GFP) in adipocytes and the ovary (**Figure 1E and F**; **Figure 3D and E**; **Figure 4A and B; Figure S7A and C**), suggesting undescribed adult specific roles for storage proteins in adult *Drosophila* females.

Consistent with our study focused on the roles of storage proteins in adult females, there is evidence that storage protein expression in adult insects is regulated in a sex-specific manner. Synthesis of both *Lsp1* and *Lsp2* occurs in adult females after transplantation of 2^nd^-instar larval fat bodies, whereas only Lsp1 synthesis can occur after transplantation in males (Butterworth et al., 1979; Jowett and Postlethwait, 1981). In *Bombyx mori*, females accumulate more storage protein compared to males (Tojo et al., 1980), and multiple insects utilize storage proteins to support egg development during adulthood (Wheeler and Buck, 1995, 1996). It has also been shown that females transcribe different hexamerin subunits compared to males (Martins et al., 2010; Tojo et al., 1980; Zakharkin et al., 2001), suggesting a female-specific systemic factor may regulate storage protein synthesis during adulthood. One possible candidate is the steroid hormone ecdysone, considering transcription of *LSPs* and *Fbp1* are ecdysone induced (Burmester et al., 1999). In *Diptera* adults, ecdysone is primarily produced by ovarian follicle cells, accumulating high levels in females but not males (Birnbaum et al., 1984; Bownes et al., 1984; Ono et al., 2006). In *Drosophila* females, increased ecdysone levels and ecdysone signaling in the nervous system promotes female feeding and the nutrient storage required for oogenesis (Sieber and Spradling, 2015). Given that formation of storage granules in larvae corresponds with an increase in ecdysone titers, female-specific activation of ecdysone signaling may provide a feed-forward mechanism for continuous storage protein production to support oogenesis and the developing egg.

### Potential roles for storage proteins during oogenesis

The presence of elevated storage proteins in adult female insects also suggests specific functions during oogenesis. Adult females produce large quantities of vitellogenin to supply developing oocytes with a surplus of amino acids to support embryonic development (Berg et al., 2024). To support developing oocytes, *Lepidoptera* adults rely on larval-derived protein reserves to promote egg production, whereas autogenous insects (such as mosquitos) store glycogen, fat, and protein in the larval fat body that is carried into adulthood (Wheeler and Buck, 1996). In contrast, the monarch butterfly produces two methionine-rich hexamerins specifically during adulthood to support egg development (Pan and Telfer, 1996). Other insects have been shown to synthesize and accumulate hexamerins, with storage protein accumulation greater in females that is quickly depleted upon vitellogenesis. For example, virgin queen ants store large quantities of storage proteins that are quickly depleted during the rearing of the first worker ants (Wheeler and Buck, 1995). Although storage proteins have been considered indispensable during *Drosophila* adulthood in some studies, our results are consistent with another report showing other adipocyte-derived factors that are primarily known for their roles in larvae also act as regulators of adult oogenesis (Simmons et al., 2025). Furthermore, we found that Lsp2::eGFP is present in the oocyte of developing egg chambers (**Figure S7D**) and both Fbp1::eGFP and Fbp2::eGFP are present in ovarian follicle cells (**Figure 4D and E**), suggesting additional roles during oogenesis [e.g., possibly mediating the uptake of Lsp2 or additional adipocyte-derived yolk proteins into the oocyte (Berg et al., 2024)] or embryonic development undiscovered by our analysis.

### Adipocyte-derived factors are required for oogenesis

Although present in other tissues, the primary site of hexamerin synthesis is in the insect fat body (Burmester and Scheller, 1997; Haunerland, 1996; Telfer and Kunkel, 1991). The fat body serves as a major endocrine organ in insects (Li et al., 2019) and fat body-specific pathways indirectly influence oogenesis, including GSC maintenance (Armstrong and Drummond-Barbosa, 2018; Armstrong et al., 2014; Matsuoka et al., 2017; Weaver and Drummond-Barbosa, 2018, 2019). For example, adipocyte-secreted collagen IV is required to maintain ovarian GSCs by promoting E-Cadherin mediated adhesion of GSCs to the niche (Weaver and Drummond-Barbosa, 2018). In addition, multiple diet-dependent pathways that regulate amino acid production are altered when adult *Drosophila* females are fed a protein-poor diet (Matsuoka et al., 2017). Furthermore, amino acid sensing in adult female adipocytes influences GSC number in a TOR-independent manner, whereas adipocyte-specific TOR activity influences ovulation (Armstrong et al., 2014). Considering storage protein synthesis in larval adipocytes occurs downstream of TOR (Valzania et al., 2024), it was interesting we did not observe defects in ovulation for when *Lsp1α*/*β*/*γ* or *Lsp2* were knocked down in adipocytes compared to control (0% of ovaries has 2 or more stage 14 embryos per ovariole for all genotypes). Therefore, our results suggest production and secretion of adipocyte-derived storage proteins may occur in a TOR-independent manner [possibly downstream of fat-specific insulin or amino acid response pathways, which also regulate GSC number (Armstrong and Drummond-Barbosa, 2018; Armstrong et al., 2014)]. Furthermore, it is possible that storage proteins act directly downstream of Svp or ecdysone signaling in adult female fat. Interestingly, publicly available data from modENCODE (Consortium, 2012; Roy et al., 2010) shows Svp binding on each storage protein loci in pupae (**Fig. S10**). In contrast, EcR in does not show high occupancy at this stage. Future studies should address whether Svp or EcR bind to regulatory regions of storage proteins to regulate their expression specifically in adult female adipocytes.

### Maintenance of GSCs requires activation of amino acid-dependent pathways

Stem cells and cancer stem cells rapidly proliferate and require increased amino acid metabolism to maintain their self-proliferative and differentiation activities (Gong et al., 2025). In *Drosophila*, larval neuroblasts require diet-derived amino acids to support growth and proliferation (Chell and Brand, 2010). Loss of the amino acid transporter *slimfast* in hematopoietic progenitors causes premature differentiation, resulting in decreased progenitor cell number (Shim et al., 2012). Storage proteins have recently been shown to accumulate in *Drosophila* larval tumors where they are required to promote cancer progression (Valzania et al., 2025). In addition, dietary methionine is required for activation of S-adenosylmethionine to regulate the division rate of intestinal stem cells (Obata et al., 2018). Adult female GSCs directly sense amino acids and activate TOR signaling to control GSC number (LaFever et al., 2010).

Our results suggest that adipocyte-derived amino acid storage proteins are required to promote TOR signaling in GSCs (**Figure 5**), suggesting that storage proteins could be trafficked to the ovary where they are catabolized to their amino acid precursors before transportation into GSCs. In larval adipocytes, Fbp1 is required for storage protein reabsorption into the fat prior to metamorphosis (Valzania et al., 2024). However, we did not observe Fbp1::eGFP localization in the GSC niche area suggesting that Fbp1 may not be mediating trafficking of storage proteins into GSCs. It is possible that Fbp1 (or Fbp2) are required in a separate tissue or cell type to catabolize LSPs to regulate GSC maintenance. Single cell RNA sequencing analysis show that *Fbp2* transcripts are present in adult gut enterocytes (Li et al., 2022). Therefore, future studies should systematically knock down *Fbp1* and *Fbp2* in different adult tissues to determine which cell type may be required for LSP absorption to control GSC number. Furthermore, multiple amino acid transporter transcripts are expressed in the ovary (Wright and Armstrong, 2024), and single cell sequencing analysis suggests that multiple transporters are expressed in female GSCs (Li et al., 2022). Future studies should also address the germline requirement for transporters that traffic aromatic amino acids and their relationship with adipocyte-derived storage proteins in regulating TOR signaling and GSC maintenance in the adult female ovary.

## DATA AVAILABILITY

*Drosophila* strains are available from the Weaver lab upon request. However, any strains that are available from the Bloomington *Drosophila* Stock Center (BDSC) must be obtained through the BDSC. All data generated for this study are included in the main text and figures or in the supplemental file provided.

## ACKNOWLEDGEMENTS

We thank the Bloomington *Drosophila* Stock Center [National Institutes of Health (NIH) P40OD018537] and the Vienna *Drosophila* Resource Center for *Drosophila* stocks. We are thankful to Flybase (www.flybase.org/), an essential *Drosophila* research resource (NIH 5U41HG000739). The authors would like to thank the Indiana University Light Microscopy Imaging Center (LMIC) for access to microscopy facilities. We are grateful to Deepika Vasudevan and Andrew Zelhof for critical reading of the manuscript. This work was supported by the NIH grants R00GM127605 and R35GM150517 to L.N.W.

## AUTHOR CONTRIBUTIONS

L.N.W. designed experiments, analyzed and interpreted data, and wrote the manuscript. A.B.Z., M.O.H., M.G.A., E.B.G., R.C.E., and L.N.W. performed experiments.

