## Supplementary Material for "Adipocyte-Derived Amino Acid Storage Proteins are Required for Germline Stem Cell Maintenance in Adult *Drosophila* Females"

**This file includes:**

Figs. S1 to S9

Table S1

### SUPPLEMENTARY FIGURES

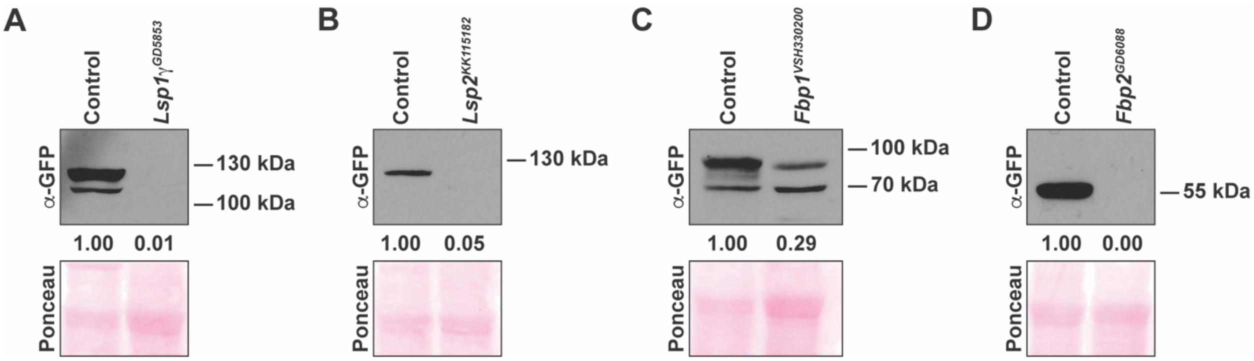


**Figure S1**. **Validation of eGFP CRISPR knock-in amino acid storage constructs.** Western blots of eGFP-tagged CRISPR knock-in constructs in control and storage protein RNAi from 3$^{rd}$ instar larvae where RNAi was induced during development using the fat body *Cg-Gal4* driver. Whole 3$^{rd}$ instar larvae were homogenized and 30 µg of total protein was loaded and probed for anti-GFP antibody for Lsp1γ::eGFP (A), Lsp2::eGFP (B), Fbp1::eGFP, and Fbp2::eGFP. Ponceau was used as the loading control.

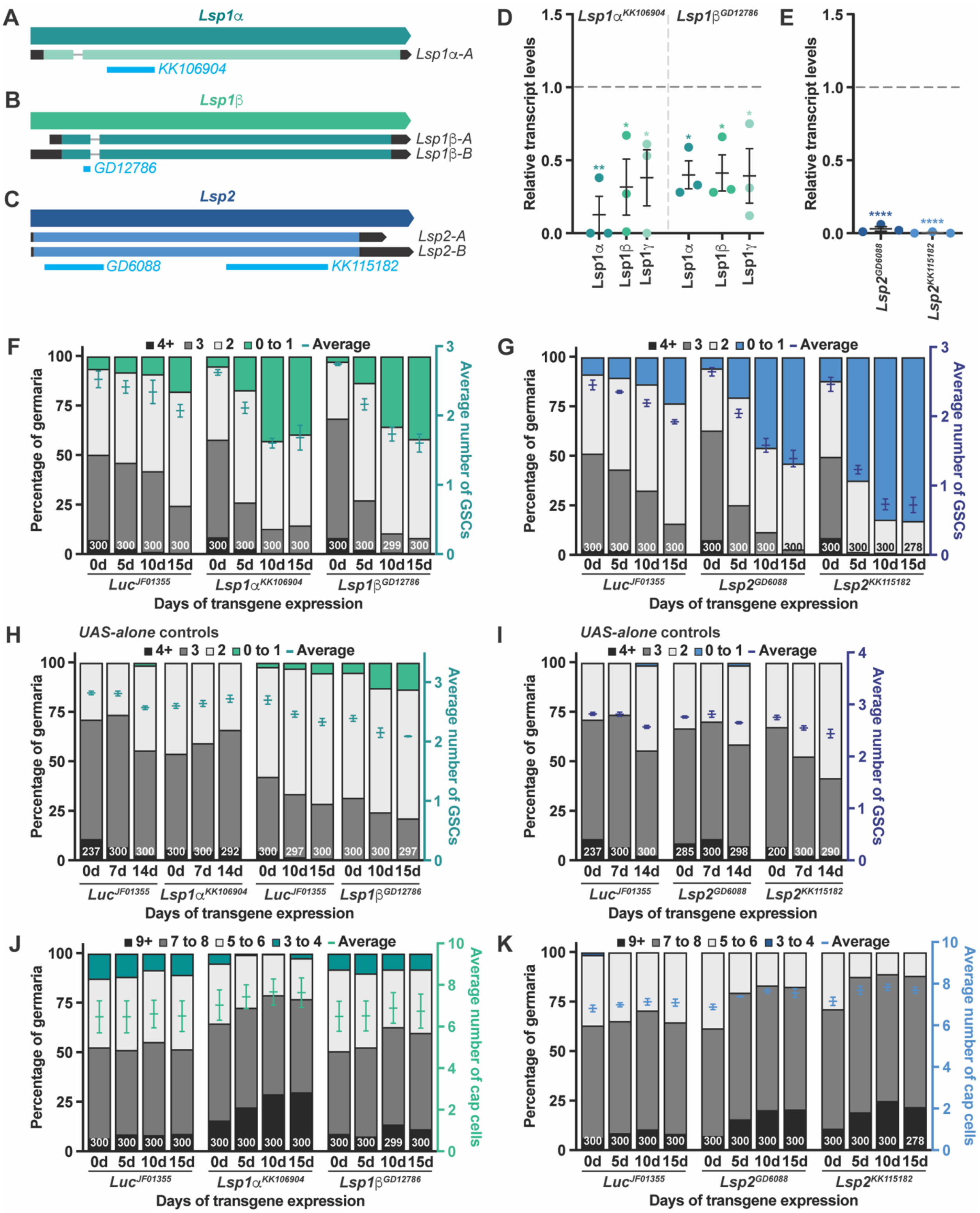
A
Lsp1α
Lsp1α-A
KK106904
B
Lsp1β
Lsp1β-A
Lsp1β-B
GD12786
C
Lsp2
Lsp2-A
Lsp2-B
GD6088
KK115182
D
Lsp1α^KK106904
Lsp1β^GD12786
Relative transcript levels
Lsp1α
Lsp1β
Lsp1γ
E
Lsp2^GD6088
Lsp2^KK115182
F
4+ 3 2 0 to 1 Average
Percentage of germaria
Average number of GSCs
0d 5d 10d 15d
Luc^JF01355
Lsp1α^KK106904
Lsp1β^GD12786
Days of transgene expression
G
4+ 3 2 0 to 1 Average
Lsp2^GD6088
Lsp2^KK115182
H
UAS-alone controls
0d 7d 14d
0d 10d 15d
I
UAS-alone controls
J
9+ 7 to 8 5 to 6 3 to 4 Average
Average number of cap cells
K
9+ 7 to 8 5 to 6 3 to 4 Average

**Figure S2. Amino acid storage proteins are required in adult female adipocytes for GSC maintenance.**

**(A-C)** Schematic showing the *Lsp1*$\alpha$ (A, teal), *Lsp1*$\beta$ (B, green), *Lsp2* (C, blue) genes and their isoforms. The RNAi hairpin lines (blue) *UAS- Lsp1*$\alpha^{KK106904}$, *UAS-Lsp1*$\beta^{GD12786}$, *UAS-Lsp2*$^{GD6088}$, and *UAS-Lsp2*$^{KK115182}$ target distinct regions of the coding regions for each transcript. **(D)** RT-qPCR analysis of *Lsp1*$\alpha$, *Lsp1*$\beta$, and *Lsp1*$\gamma$ transcripts from isolated adult female fat bodies at 10 days of adipocyte-specific knockdown against *Lsp1*$\alpha^{KK106904}$ and *UAS-Lsp1*$\beta^{GD12786}$ relative to *Luc* control using the *3.1Lsp2*$^{ts}$ driver. Knockdown of *Lsp1*$\alpha$ or *Lsp1*$\beta$ is sufficient to simultaneously decrease transcript levels of *Lsp1*$\alpha$, *Lsp1*$\beta$ and *Lsp1*$\gamma$. Data shown as mean ± standard error of the mean (SEM) from three independent experiments. *$P$<0.05, **$P$<0.01; Student's *t*-test. **(E)** RT-qPCR analysis of *Lsp2* transcripts from isolated adult female fat bodies after 10 days of adipocyte-specific knockdown relative to *Luc* control. Data shown as mean ± standard error of the mean (SEM) from three independent experiments. ****$P$<0.0001; Student's *t*-test. **(F,G)** Bar graphs representing the percentage of germaria containing zero-to-one, two, three, or four-or-more GSCs at different days of adult adipocyte-specific RNAi against *Luc* control, *Lsp1*$\alpha$, *Lsp1*$\beta$, or *Lsp2* (left y-axis). GSC number averages shown as mean ± standard error of the mean (SEM; right y-axis) are also plotted in **Figure 2C and D**. Numbers of germaria analyzed are shown inside bars. **(H,I)** Data represented as in (F) and (G) for *UAS* alone control females carrying *UAS-hairpin* transgenes against *Lsp1*$\alpha/\beta$ (H) or *Lsp2* (I) relative to *Luc* control in the absence of *3.1Lsp2*$^{ts}$ and subjected to similar temperature shift as in (F,G). No statistical differences from three biological replicates, two-way ANOVA with interaction. The number of germaria analyzed are shown in each bar. **(J,K)** Bars representing the percentage of germaria containing three-to-four, five-to-six, seven-to-eight, or greater or equal to nine cap cells per germarium (left y-axis) at different times of adult adipocyte-specific knockdown of *Luc* control, *Lsp1*$\alpha$, *Lsp1*$\beta$, or *Lsp2* (left y-axis). Average numbers of cap cells per germarium shown as mean ± standard error of the mean (SEM; right y-axis). Numbers of germaria analyzed are shown inside bars. No statistical differences from three biological replicates, two-way ANOVA with interaction.

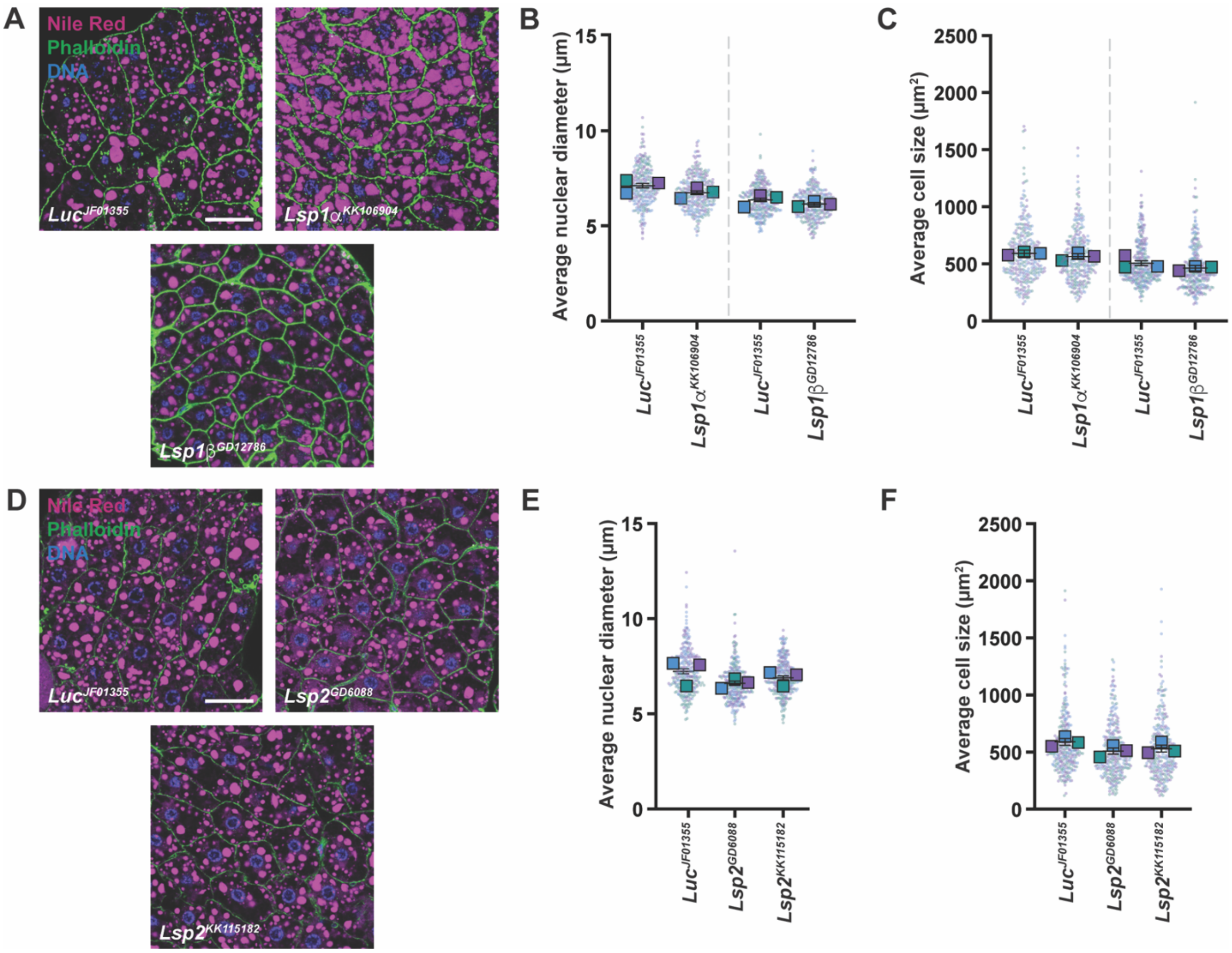


**Figure S3. Adipocyte morphology is not altered when storage proteins are knocked down in adult female adipocytes.**

**(A)** Adipocytes from females at 10 days of adult-specific RNAi against *Luc* control, *Lsp1*$\alpha$, or *Lsp1*$\beta$ in adipocytes and analyzed for adipocyte morphology. Nile Red (magenta), lipid droplets; Phalloidin (green), cell membrane; DAPI (blue), nuclei. Scale bar = 25 µm. **(B,C)** The average cell area (**B**) or average nuclear diameter of adipocyte nuclei (**C**). The averages from three independent experiments are shown as mean ± 95% confidence interval. 300 adipocytes or nuclei were analyzed. No statistically significant differences, Student's *t*-test. **(D-F)** Fat body images (**D**), average adipocyte nuclear diameter (**E**), and average adipocyte cell size (**F**) of females from *Luc* control or *Lsp2* adipocyte-specific knockdown under similar conditions as in

**A-C**. 300 adipocytes or nuclei were analyzed from three independent experiments. No statistically significant differences; Student’s *t*-test.

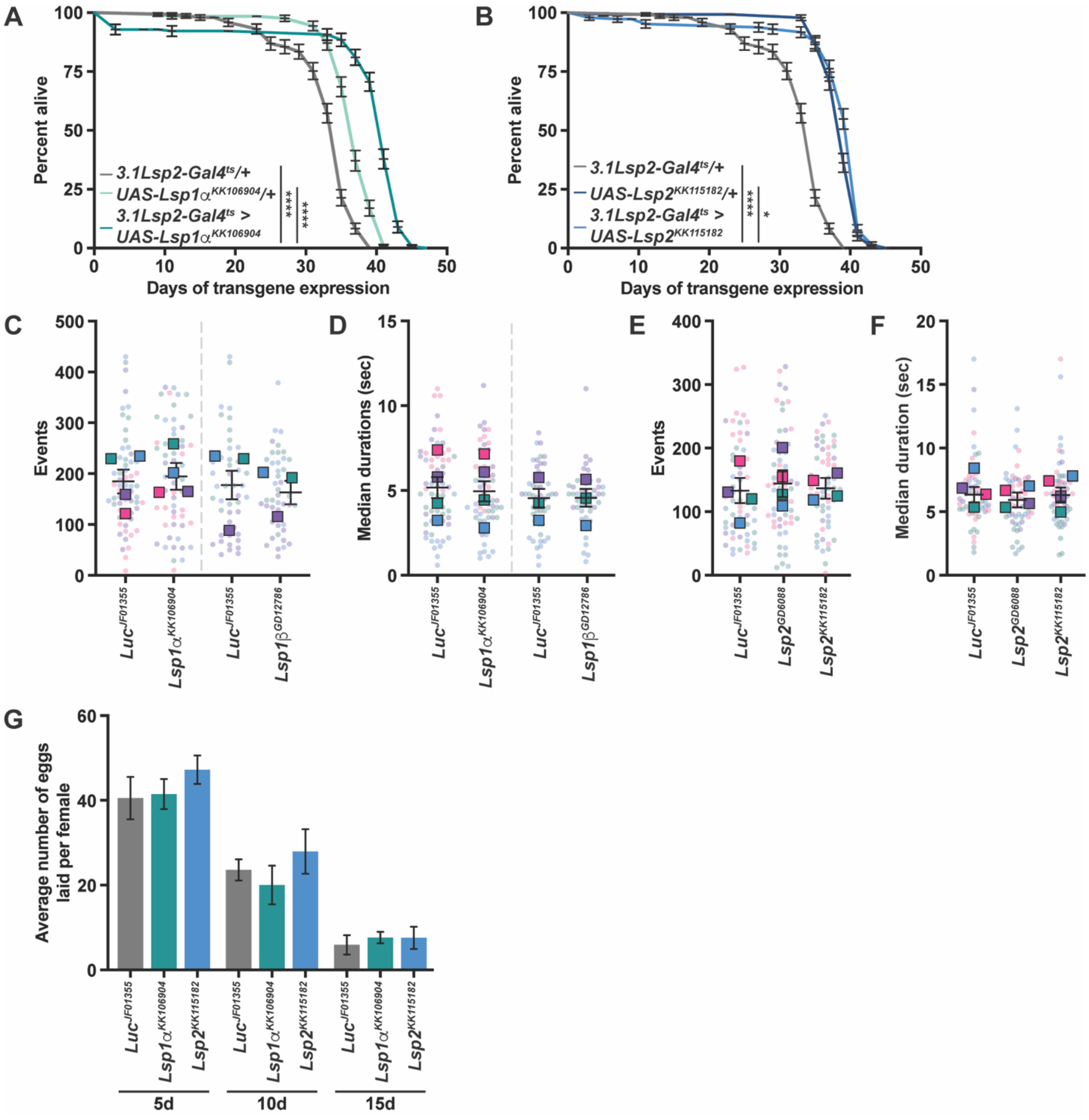


**Figure S4. Adipocyte-specific loss of storage proteins in adult females increases lifespan but does not influence feeding behavior or egg production.**

**(A,B)** Lifespan of adult females with adipocyte specific knockdown of *Lsp1α* (A) or *Lsp2* (B) compared to *UAS-alone* and *3.1Lsp2-Gal4$^{ts}$-alone* controls. At least 120 females per condition were analyzed. **P*<0.05, *****P*<0.0001; log-rank test. **(C–F)** The total number of feeding events and median time of feeding activity from adult females with adipocyte-specific knockdown of

*Lsp1α/β* (**C,D**) or *Lsp2* (**E,F**) after 10 days of transgene induction relative to *Luc* control. Data shown as mean ± 95% confidence interval. At least 50 females were analyzed from three independent experiments. No statistically significant differences, Mann-Whitney *U*-test. **(G)** The average number of eggs laid by females with adult adipocyte-specific knockdown of *Lsp1a* or *Lsp2* relative to *Luc* control over time. Data shown as mean ± SEM. No statistical differences.

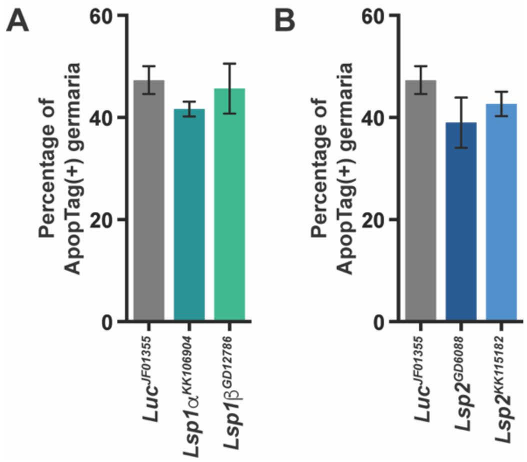


**Figure S5. Adipocyte-specific knockdown of storage proteins does not cause germline cyst death.**

**(A,B)** Percentage of germaria containing ApopTag-positive cysts (indicative of cell death) in adult females from adipocyte-specific knockdown *Lsp1α/β* RNAi (A) or *Lsp2* RNAi (B) relative to *Luc* control using the *3.1Lsp2-Gal4*$^{ts}$ driver. No significant differences between storage protein knockdown and control from three independent experiments, Student's *t*-test.

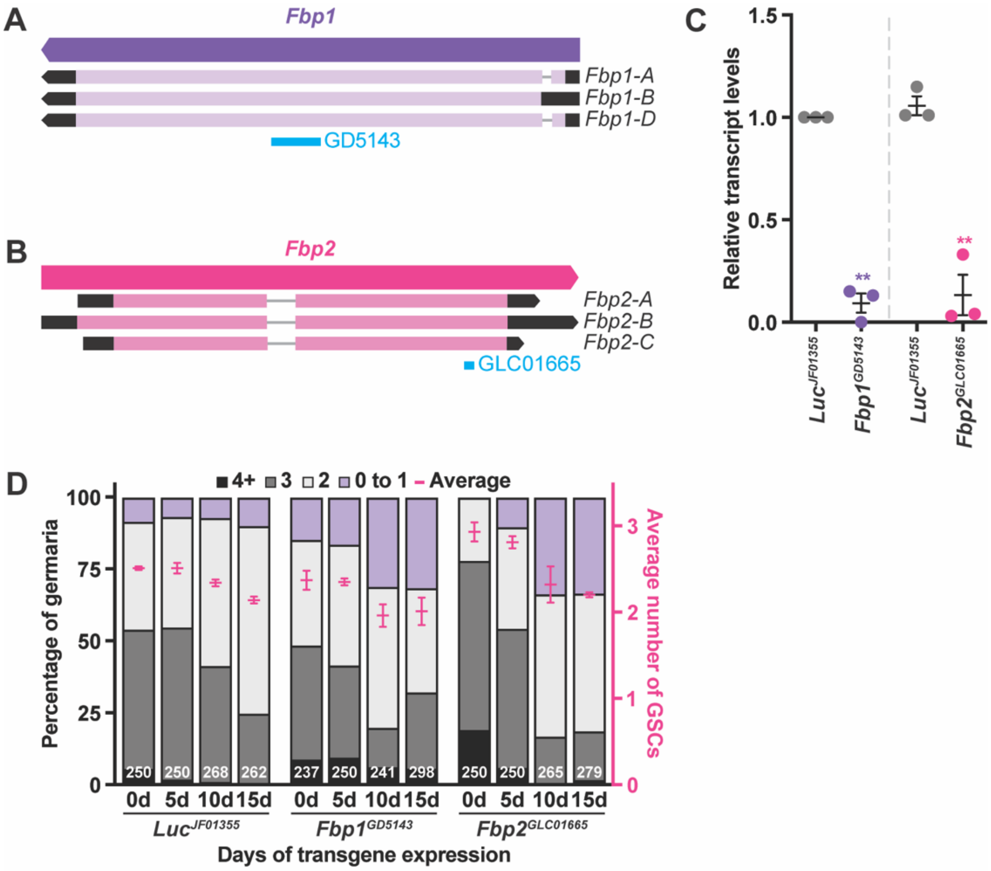


**Figure S6. Fbp1 nor Fbp2 are required in adult female adipocytes for GSC number.** **(A,B)** Schematic showing the *Fbp1* (**A**, purple) and *Fbp2* (**B**, pink) genes and their three isoforms. The RNAi hairpin lines (blue) *UAS-Fbp1*$^{GD5143}$ and *UAS-Fbp2*$^{HMS02769}$ target all isoforms of the respective gene. **(C)** RT-qPCR analysis of *Fbp1* and *Fbp2* transcripts from isolated adult female fat bodies at 10 days of adipocyte-specific knockdown against *Fbp1* or *Fbp2* relative to *Luc* control using the *3.1Lsp2*$^{ts}$ driver. Data shown as mean ± SEM from three independent experiments. ***P*<0.01; Student's *t*-test. **(D)** Bar graph representing the percentage of germaria containing zero-to-one, two, three, or four-or-more GSCs at different days of adult adipocyte-specific RNAi against *Fbp1* or *Fbp2* relative to *Luc* control (left y-axis). GSC number averages shown as mean ± standard error of the mean (SEM; right y-axis) are also plotted in **Figure 3C**. The number of analyzed germaria are shown in the bars.

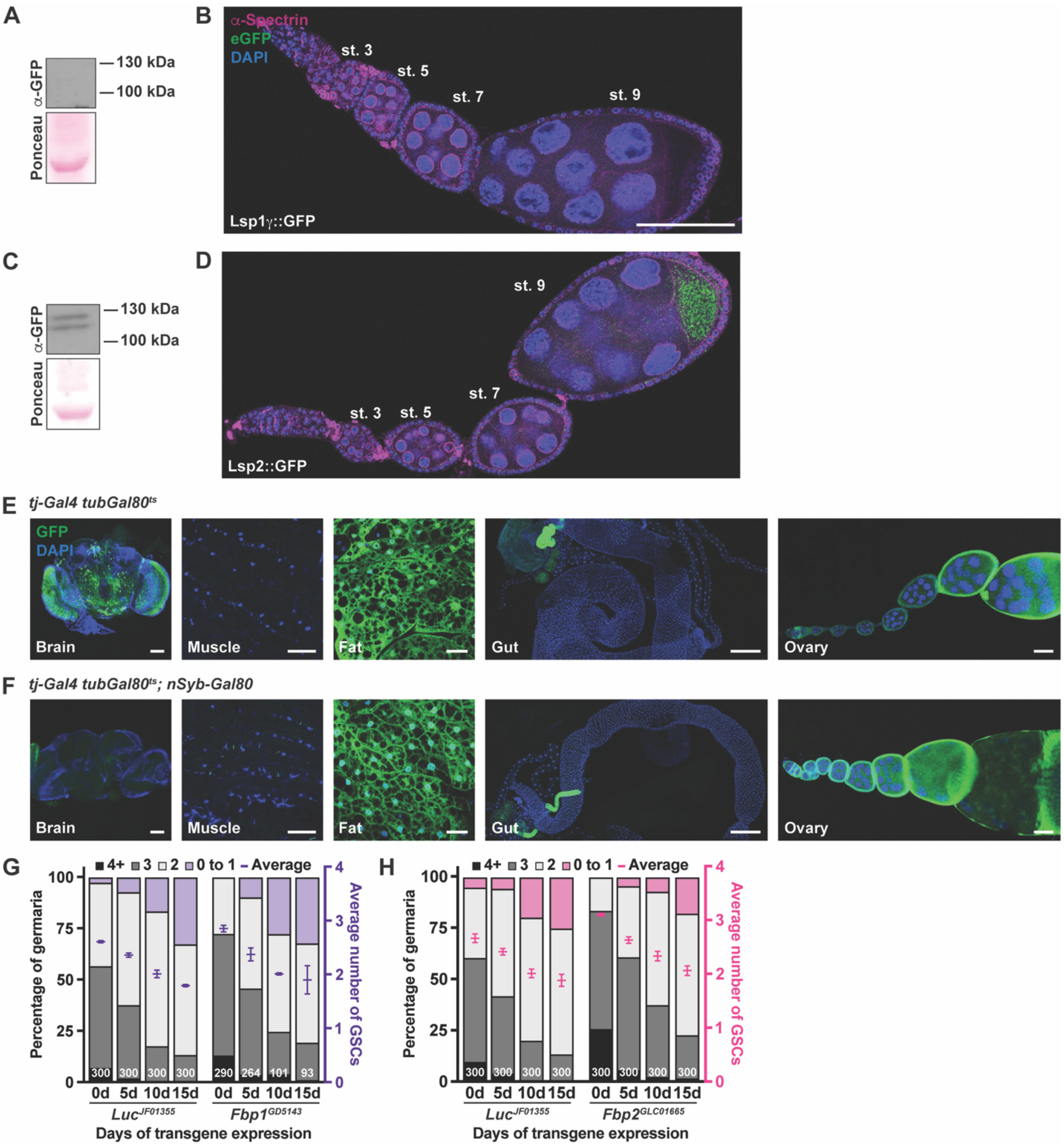


**Figure S7. Fbp1 and Fbp2 are expressed in ovaries, but adult follicle cell knockdown of *Fbp1* nor *Fbp2* decreases GSC number.**

**(A,C)** Western blots from adult female ovaries in which 30 µg of total protein was loaded and probed with anti-GFP antibodies to detect Fbp1::eGFP (A) or Fbp2::eGFP (C). Ponceau was used as the loading control. **(B,D)** Ovarioles from adult females expressing CRISPR knock-in eGFP::fusions proteins for Lsp1γ (B, representing the Lsp1 complex) and Lsp2 (D). eGFP (green), amino acid storage fusion protein; α-spectrin (magenta), fusome; LamC (magenta), cap cell nuclear lamina; DAPI (blue), nuclei. Scale bar = 100 µm. **(E,F)** Representative images of adult female brains, muscle, fat, guts, and ovarioles from *tj-Gal4 tubGal80*$^{ts}$ (E) or *tj-Gal4 tubGal80*$^{ts}$*; nSyb-Gal80* (F) females expressing *UAS-nucGFP*. GFP (green), tissues/cells that express the *Gal4* transgene; DAPI (blue), nuclei. Scale bars = 100 µm (brain), 50 µm (muscle), 25 µm (fat), 250 µm (gut), and 100 µm (ovary). (G,H) Bar graphs representing the percentage of germaria containing zero-to-one, two, three, or four-or-more GSCs at different days of adult adipocyte-specific RNAi against *Luc* control, *Fbp1* (G) or *Fbp2* (H) (left y-axis). GSC number averages shown as mean ± standard error of the mean (SEM; right y-axis) are also plotted in **Figure 4F and G**. Numbers of germaria analyzed are shown inside bars.

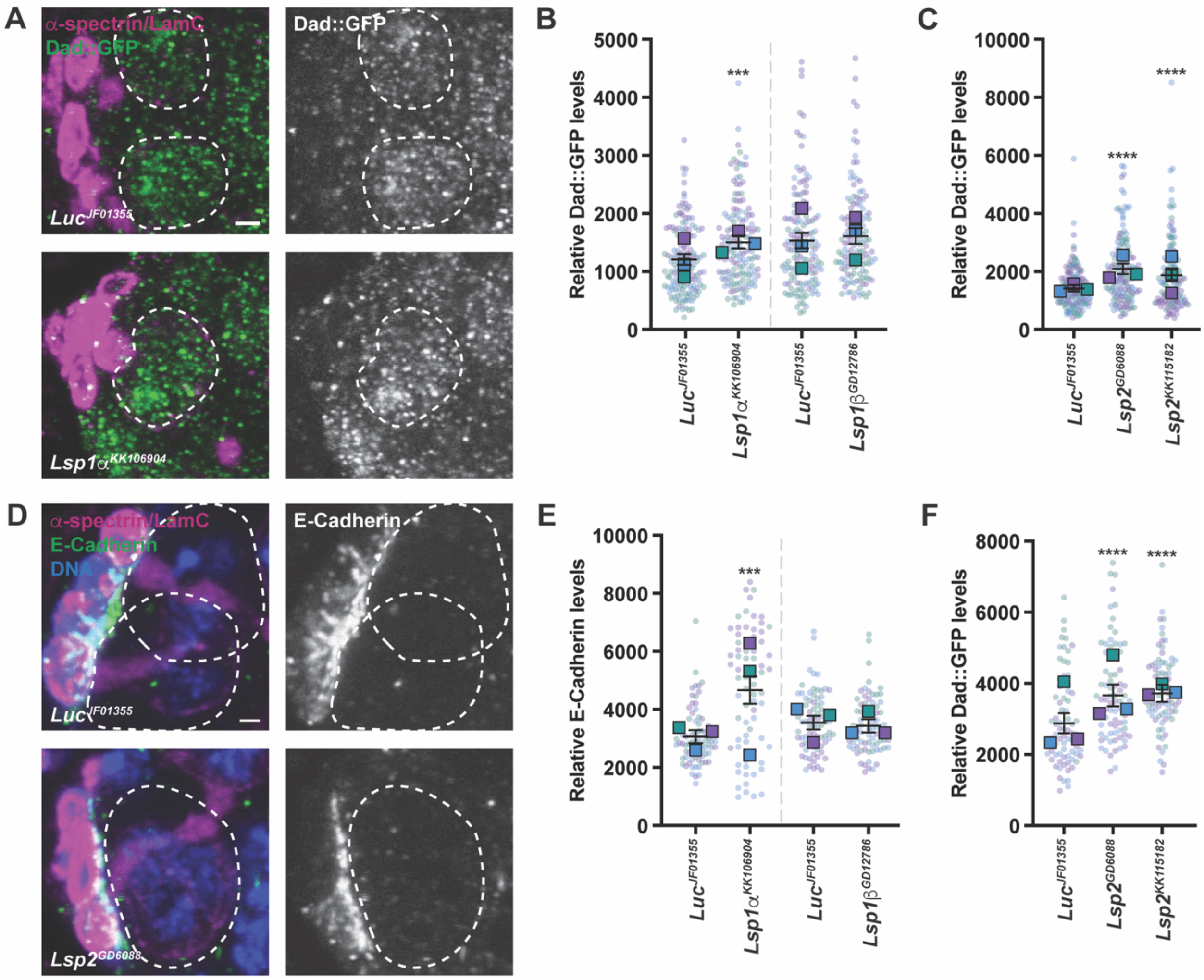


**Figure S8. Adipocyte-specific knockdown of amino acid storage proteins does not decrease BMP signaling or E-Cadherin levels in the ovary.**

**(A)** Anterior portion of the germaria from females after 10 days of adult-specific knockdown of *Luc* control or *Lsp1α* in adipocytes using the *3.1Lsp2^ts* driver showing similar levels of nuclear Dad [Dad::nlsGFP; a reporter of BMP signaling (Ayyaz et al., 2015)]. Dad::nlsGFP (green); α-spectrin (magenta), fusome; LamC (magenta), cap cell nuclear lamina. GSC nuclei are outlined. Scale bar = 2.5 µm. **(B,C)** Superplots of mean Dad::nlsGFP intensity per GSC in experiment in (A) for *Lsp1α/β* (B) or *Lsp2* (C) knockdown in adipocytes. Data shown as mean ± 95% confidence interval. ****P* < 0.001, *****P* < 0.0001; Mann Whitney *U*-test. At least 125 GSCs were analyzed for each genotype from three independent experiments. **(D)** Anterior portion of the germaria from females at 10 days of adult adipocyte-specific *Luc* control or *Lsp2* RNAi showing similar levels of E-Cadherin at the GSC-niche junction. E-Cadherin (green); $\alpha$-spectrin (magenta), fusome; LamC (magenta), cap cell nuclear lamina. GSCs are outlined. Scale bar = 2.5 µm. **(E,F)** SuperPlots of total GSC-niche junction E-Cadherin intensity per germarium for the experiment in (**D**) for adipocyte-specific knockdown of *Lsp1*$\alpha/\beta$ (E) or *Lsp2* (F). Data shown as mean ± 95% confidence interval. At least 75 germaria were analyzed for each condition. ***$P$ < 0.001, ****$P$ < 0.0001; Mann Whitney $U$-test.

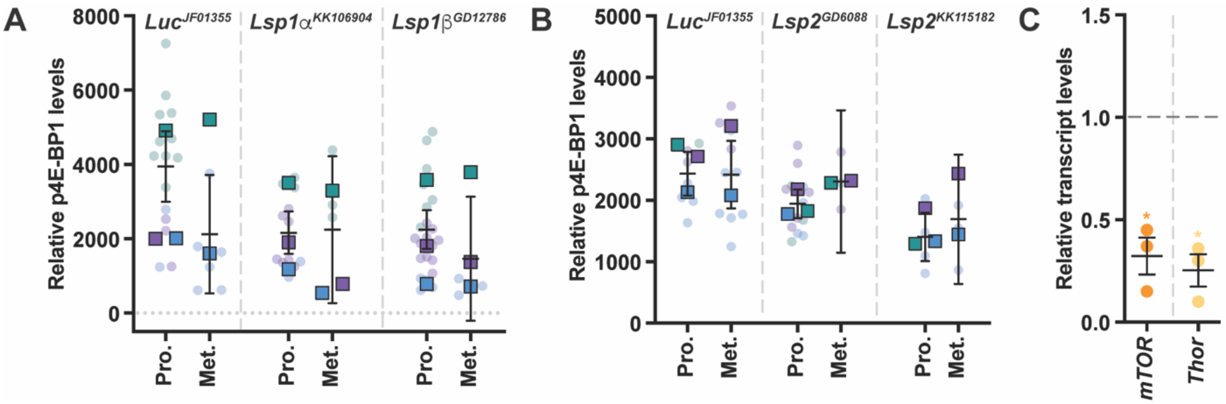


**Figure S9. p4E-BP1 levels in GSCs are similar during early mitotic stages.**

**(A,B)** SuperPlots showing p4E-BP1 levels in GSCs during prometaphase (Pro) or metaphase (Met) for adipocyte-specific knockdown of *Lsp1α* (A) or *Lsp2* (B) compared to *Luc* controls. Data are shown as mean ± 95% confidence interval and is also shown in **Figure 5B**. **(C)** RT-qPCR of transcript levels for *mTOR* or *Thor* in *mTOR*$^{\Delta P}$*/CyO; 3.1Lsp2-Gal4*$^{ts}$ or *Thor*$^{2}$*/CyO; 3.1Lsp2-Gal4*$^{ts}$ females, respectively.

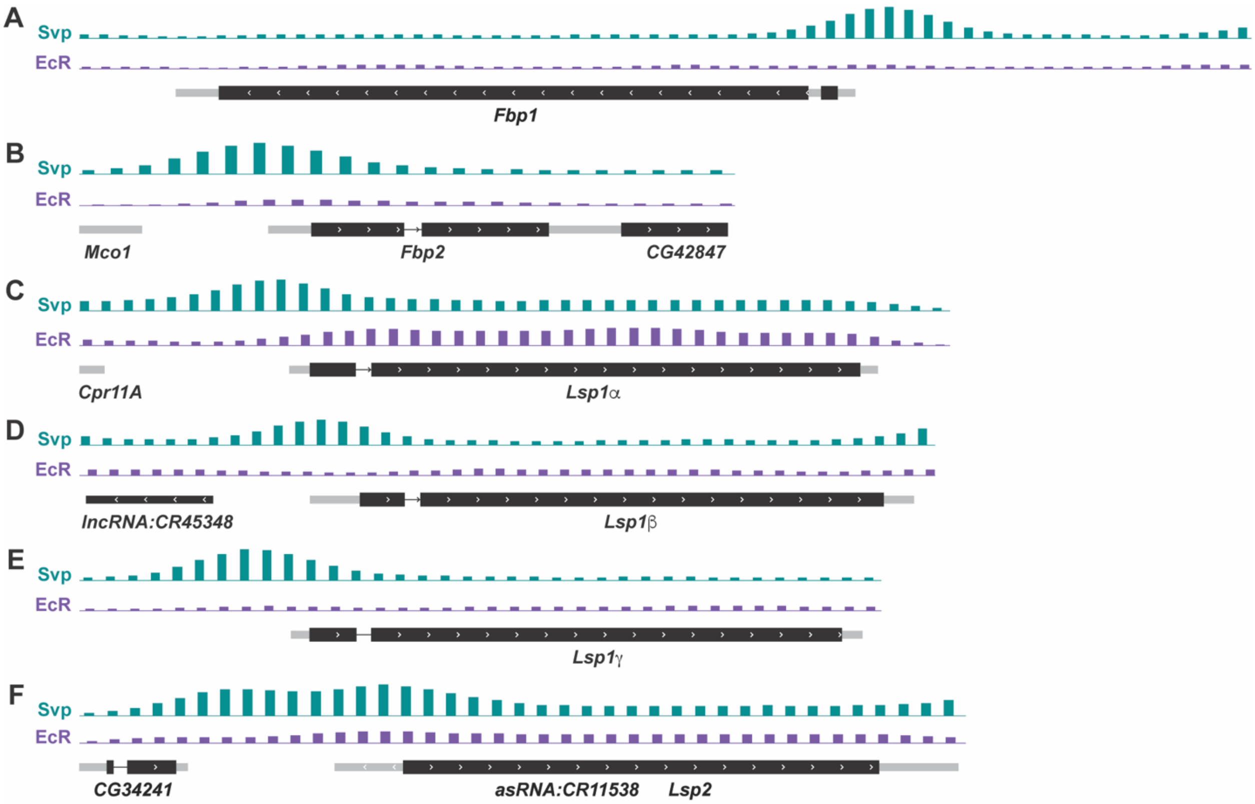


**Figure S10. Amino acid storage proteins contain Svp binding sites.**

Analysis of moENCODE data occupancy of Svp and EcR on the genetic loci of *Fbp1* (A), *Fbp2* (B), *Lsp1*$\alpha$ (C), *Lsp1*$\beta$ (D), *Lsp1*$\gamma$ (C), and *Lsp2* (D).

### SUPPLEMENTARY TABLES

**Table S1. Primer sequences used in this study.**

| Gene | Forward | Reverse |
|---|---|---|
| ***Lsp1*$\alpha$** | 5’-ACATCAAGGTCGCTGACAAGG-3’ | 5’-TTCCCATCGCAATGTACTCTTC-3’ |
| ***Lsp1*$\beta$** | 5’-GATCGCCATCGCATTGCTG-3’ | 5’-CCCTGCTTGATGTGGTCCT-3’ |
| ***Lsp1*$\gamma$** | 5’-GCCTGTGTGACTGCCTTTAG-3’ | 5’-AGAGGCTCATCAATACGGTGA-3’ |
| ***Lsp2*** | 5’-CTTCCAGCACGTCGTCTACTG-3’ | 5’-CCCTGCATATCATCACGGAACA-3’ |
| ***Fbp1*** | 5’-CTTCGCCGTAATGTGGTCTAC-3’ | 5’-AGAGCTTGAGTGTCCTCACGA-3’ |
| ***Fbp2*** | 5’-ATGAATCTGACTGGCATGATCCA-3’ | 5’-CCAGGCCATAGACAGAGGACA-3’ |
| ***mTOR*** | 5’-CAGATGCCCGAGGTGTACTC-3’ | 5’-CATGAAAGCCCGCTCGTAGA-3’ |
| ***Thor*** | 5’-TTACTGCCAGGAAGGGCATTT-3’ | 5’-TGACGGACACATCGTTGATTA-3’ |
| ***Rp49*** | 5’-CAGTCGGATCGATATGCTAAGC-3’ | 5’-AATCTCCTTGCGCTTCTTGG-3’ |
| ***Act5c*** | 5’- AGGCCAACCGTGAGAAGATG-3’ | 5’- ACATACATGGCGGGTGTGTT-3’ |
